# How important is it to consider lineage diversification heterogeneity in macroevolutionary studies: lessons from the lizard family Liolaemidae

**DOI:** 10.1101/563635

**Authors:** Olave Melisa, Luciano J. Avila, Jack W. Sites, Morando Mariana

**Affiliations:** Department of Biology, University of Konstanz, Konstanz, Germany; Instituto Argentino de Investigaciones de Zonas Arídas, Consejo Nacional de Investigaciones Cientificas y Técnicas (IADIZA-CONICET), 5500 Mendoza, Argentina; Instituto Patagónico para el Estudio de los Ecosistemas Continentales, Consejo Nacional de Investigaciones Científicas y Técnicas (IPEEC-CONICET), Boulevard Almirante Brown 2915, U9120ACD, Puerto Madryn, Chubut, Argentina; Department of Biology and M.L. Bean Life Science Museum, Brigham Young University (BYU), Provo, UT 84602, USA; Department of Biology, Austin Peay State University, Clarksville, Tennessee, 37044; Universidad Nacional de la Patagonia San Juan Bosco, Sede Puerto Madryn, Boulevard Almirante Brown 3700, U9120ACD, Puerto Madryn, Chubut, Argentina

**Keywords:** GeoHiSSE, GeoSSE, hidden states, RPANDA, speciation, extinction, macroevolution, biogeography, Andes

## Abstract

Macroevolutionary and biogeography studies commonly apply multiple models to test state-dependent diversification. These models track the association between states of interest along a phylogeny, but many of them do not consider whether different clades might be evolving under different evolutionary drivers. Yet, they are still commonly applied to empirical studies without careful consideration of possible lineage diversification heterogeneity along the phylogenetic tree. A recent biogeographic study has suggested that orogenic uplift of the southern Andes has acted as a species pump, driving diversification of the lizard family Liolaemidae (307 described species), native to temperate southern South America. Here, we argue against the main findings of this study that attributes the Andean uplift as main driver of the evolution of this group. We show that there is a clear pattern of heterogeneous diversification in the Liolaemidae, which biased the state- and environment-dependent analyses performed in the GeoSSE and RPANDA programs, respectively. We show here that there are two shifts to accelerated speciation rates involving two clades that were both classified as having “Andean” distributions. We incorporated the Geographic Hidden-State Speciation and Extinction model (GeoHiSSE) to accommodate unrelated diversification shifts, and also re-analyzed the data in RPANDA program after splitting biologically distinct clades for separate analyses, as well as including a more appropriate set of models. We demonstrate that the “Andean uplift” hypothesis is not supported when the heterogeneous diversification histories among these lizards is considered. We use the Liolaemidae as an ideal system to demonstrate potential risks of ignoring clade-specific differences in diversification patterns in macroevolutionary studies. We also implemented simulations to show that, in agreement with previous findings, the HiSSE approach can effectively and substantially reduce the level of distribution-dependent models receiving the highest AIC weights in such scenarios. However, we still find a relatively high rate of distribution-dependent models receiving the highest AIC weights (= 15%), and provide recommendations related to the set of models included in the analyses that reduce these rates by half. Finally, we demonstrate that trees including clades following different dependent-drivers affect RPANDA analyses by producing different outcomes, ranging from partially correct models to completely misleading results. We provide recommendations for the implementation of both programs.

## Introduction

Macroevolutionary modeling of diversification plays an important role in inferring large-scale biodiversity patterns (Schluter 2016). Several studies have focused on quantifying differences in macroevolutionary patterns linked to geographic, ecological, life-history and other traits, based on the variation in speciation and extinction rates (Jablonski 2008; Rabosky and McCune 2010; Condamine et al. 2013; Ng and Smith 2014). Given that mechanisms underlying the correlations between characters and diversification are generally poorly understood (Rabosky and Goldberg 2015), a family of State-Dependent Speciation and Extinction (SSE) models have been developed. The earliest models included binary traits (Maddison 2006), quantitative traits (FitzJohn 2010), geographic character states (Goldberg et al. 2011), multiple character states (FitzJohn 2012), punctuated trait changes (Goldberg and Igic 2012; Magnuson-Ford and Otto 2012), and time-dependent macroevolutionary rates (Rabosky and Glor 2010). These models track associations between the states of interest and speciation and extinction rates along a phylogenetic tree, and they have been implemented in hundreds of empirical studies (Rabosky and Goldberg 2015). In addition, interest has also increased in the application of correlative algorithms (Steeman et al. 2009; Winkler et al. 2010; Condamine et al. 2012; 2013; Morlon et al. 2016), as implemented in the software RPANDA (Morlon et al. 2016), to use environment-dependent models to test whether gradual changes in paleo-environments have significantly influenced speciation and extinction rates. These state and environment-dependent models have become popular, but important concerns have been raised (at least) for the SSE family of models that do not consider whether unrelated traits are associated with shifts in diversification rates (Maddison and FitzJohn 2014; Rabosky and Goldberg 2015; 2017; Beaulieu and O’Meara 2016). For example, strong correlations with diversification are sometimes inferred from rate shifts, leading to dramatically high rates of false positives even if the shift is unrelated to the targeted state (Maddison et al. 2007; FitzJohn 2010; Maddison and FitzJohn 2014; Rabosky and Goldberg 2015; 2017). These issues may also be affecting environment-dependent correlative models in such scenarios (Lewitus and Morlon 2018). Consequently, while larger trees are preferred due to the presumed increase in power, this also increases the risk of including clades that differ in factors that can affect diversification rates along a tree (ecological requirements, dispersal abilities, and life history traits [Li et al. 2018]). Hence, at least for the case of SSE models, new algorithms that include “hidden states” (HiSSE) have been proposed for binary traits (BiHiSSE; Beaulieu and O’Meara 2016), and more recently geographic-dependent diversification hypotheses (GeoHiSSE; Caetano et al. 2018) and multiple states (SecSSE; Herrera-Alsina et al. 2019). Hidden states refer to unsampled traits that are related to diversification shifts in a phylogenetic tree, thus incorporating heterogeneous diversification into the original SSE.

The HiSSE has improved the SSE model family and potentially resolved the problem of unrelated shifts in the phylogeny when appropriate sets of models are tested (Beaulieu and O’Meara 2016; Caetano et al. 2018; Herrera-Alsina et al. 2019). On the other hand, there are no currently available solutions for environment-dependent models (i.e. hidden states have not been implemented so far), besides splitting clades into separate analyses (Lewitus et al. 2018). Even though models that do not include hidden states are not suitable for trees including groups with heterogeneous diversification patterns, they are still commonly applied in macroevolutionary studies without careful assessment. For instance, the Geographic State Speciation and Extinction model (GeoSSE; Goldberg et al. 2011) was recently used in biogeographic studies of lizards (Esquerré et al. 2019), tanagers and tortoises (Román-Palacios and Wiens 2018), plants (Canal et al. 2019), including palms (Bacon et al. 2018) and oaks (Hipp et al. 2018), and a recent mega-phylogeny of mushrooms (Varga et al. 2019). Given the above-noted limitations of this model, these empirical cases should be revisited (e.g. Harrington and Reeder 2017). Here, we re-analyze the recently published phylogeny of the lizard family Liolaemidae (Esquerré et al. 2019), and show how the main conclusion can change when considering heterogeneous diversification in state- and environment-dependent hypotheses tests.

The lizard family Liolaemidae is the most species-rich lizard clade in southern South America (307 species; Reptile Database 11 February 2019). The clade includes three genera: *Ctenoblepharys, Liolaemus* and *Phymaturus* (Table 1). *Ctenoblepharys* is a monotypic genus restricted to the coastal desert of Peru (Table 1), whereas *Liolaemus* is the world’s richest temperate zone genus of extant amniotes (Olave et al. 2018), with 262 described species (Reptile Database 2 February 2019). This clade includes a highly diverse group of species inhabiting a wide range of different environments (Table 1), and the sister genus *Phymaturus* (44 species; Reptile Database 2 February 2019) is distributed along both the eastern and western Andean slopes in Argentina and Chile (*palluma* clade), and through Patagonia (*patagonicus* clade). *Phymaturus* are highly specialized lizards, strictly saxicolous and largely restricted to volcanic plateaus and peaks (Cei 1986).

**Table 1:** Summary of distribution, habitat use, diet and reproductive mode among the three Liolaemidae genera.

|  | *Ctenoblepharys* | *Phymaturus* | *Liolaemus* |
| --- | --- | --- | --- |
| **Described species** | 1 | 44 | 262 |
| **Distribution** | Perú | Argentina<br>Chile | Argentina<br>Chile<br>Perú<br>Bolivia<br>southern Brazil<br>Paraguay<br>Uruguay |
| **Habitat** | coastal desert | saxicolous | terrestrial<br>arboreal<br>arenicolous<br>saxicolous |
| **Diet** | insectivores | herbivores | herbivores<br>omnivores<br>insectivores |
| **Time for sexual maturity** | unknown | 7-8 years | ~2 years |
| **Reproductive mode** | oviparous | viviparous | viviparous<br>oviparous<br>parthenogenesis |

The three genera have clear differences in species richness, ecological requirements, behaviors, and life histories (Table 1). A recent macroevolutionary study found disparate patterns of diversification among the three genera (Olave et al. *in press*), while another recent biogeographic study focused on the entire clade, overlooking these striking differences between *Phymaturus* and *Liolaemus*, and the diversification rate shifts along the tree (Esquerré et al. 2019). The Esquerré et al. study represents a major contribution to evolutionary biology and herpetology in that it: (i) presents the largest Liolaemidae time-calibrated phylogeny to date (258 taxa), (ii) the most extensive compilation of habitats, altitudes, and temperature data for all taxa, and (iii) documents shifts in parity modes, including reversals from viviparity to oviparity, a transition believed to be rare. However, Esquerré et al. approach the state-dependent hypothesis using the GeoSSE to test for differences in speciation rates in Andean vs non-Andean (low elevation) species, and detected higher speciation rates in the Andean areas. They also performed correlative environment-dependent analyses in RPANDA program and found a significant exponential relationship of speciation rate with Andean uplift. Combining these results with ancestral area reconstructions, the authors infer that the Andean orogeny has acted as a “species pump” and that it is the main driver of diversification in the evolution of Liolaemidae.

Here, we show that clade-specific differences within the Liolaemidae have affected the conclusions drawn from GeoSSE and RPANDA. We re-analyzed this dataset and show that Andean uplift-dependency is not supported as the main driver of Liolaemidae diversification, when considering heterogeneous diversification across the clade’s phylogenetic history. In this study, we (i) present an alternative explanation to Esquerré et al., to describe the diversification patterns within the Liolaemidae; and (ii) based on simulation studies, suggest recommendations that should be considered in empirical studies using the state- (HiSSE) and environment-dependent (RPANDA) models.

## Materials and methods

Here we briefly describe our methods, but for further details, see the Extended Materials and Methods in the Supplementary Material.

### Identifying possible diversification shifts along the tree and exploration of the relationship between speciation rates vs. altitude

We explored the association between speciation (and extinction) rates with the maximum altitude of occurrence known for any species (phylogentic tree and altitude data taken from Esquerré et al. 2019). The phylogenetic tree includes the monotypic *Ctenoblepharys*, 188 described + 11 undescribed species of *Liolaemus*, and 35 described + 23 undescribed species of *Phymaturus* (73% species coverage of all recognized Liolaemidae). We estimated speciation and extinction rates using BAMM 2.5 (Rabosky et al. 2014) because the method estimates rates per branch, allowing us to compare changes of these rates among the clades and species (i.e., tips) of interest. We also compared BAMM results with the recently introduced ClaDS program (Maliet et al. 2019).

### Hypothesis testing: role of the Andean orogeny in diversification of the Liolaemidae

#### Geographic Hidden-State Speciation and Extinction (GeoHiSSE)

We then implemented the GeoHiSSE model (Caetano et al. [2018]) using the *hisse* package (Beaulieu and O’Meara 2016). Given that we do not fully agree with the original classification of Andean and non-Andean species by Esquerré et al. (2019), here we propose an alternative classification, and performed these analyses using both (Table S8). As an example, the “Patagonia” group is distributed across a huge area that was assumed to be “Andean” in Esquerré et al. (2019), which we consider a poor classification for many species. For example, both the *P. patagonicus* and the *L. lineomaculatus* clades are restricted mainly to the lowland Patagonian steppe. Generally, the Esquerré et al. (here after the “original classification”) classification recognizes 194 Andean species, 39 non-Andean species, and 25 widespread species; here our preferred classification includes 118 Andean species, 121 non-Andean species, and 19 widespread species.

We first evaluated the set of 35 different models proposed by Caetano et al. (2018), and performed simulations based on 200 random permutations of geographic distributions (Rabosky and Golberg 2015) to assess power (for details in simulations see Extended Materials and Methods). Permutations were performed by keeping the Liolaemidae phylogenetic tree, thus maintaining the complexity of the empirical scenario. This strategy allows us to assess the expected rate of distribution-dependent models receiving the highest AIC weights given the specific phylogeny of Liolaemidae. We performed all simulations twice, based on the original and our preferred geographical classification. The models include a wide range of possible scenarios, including 25 area-independent models (CID) indicating no relationship between diversification rates and geographic areas, and 10 area-dependent diversification models (all models listed in Table S7). Given the relatively high rates distribution-dependent models receiving the highest AIC weights found in our simulations (see results), and issues of overparametrization (Caetano et al. 2018), we removed some of the models from the original 35. Specifically, since some of the proposed state-dependent models have a greater number of free parameters than the most complex null models that *hisse* can currently implement (15 free parameters in the most complex CID model [M11 and M17, both including 5 hidden classes]), we suggest that excluding models with >15 free parameters is important to reduce these rates. Thus, we consider only 32 models, excluding the most complex models (M10, M16 and M32, with 19, 21 and 17 free parameters, respectively).

#### Correlative environment-dependent models in RPANDA

Esquerré et al. tested whether speciation and extinction rates are correlated with the Andean uplift using the R package RPANDA (Morlon et al. 2016). Similar to the description above, they implemented this model on the complete phylogeny of Liolaemidae and did not analyze the two different genera into separate analyses. They fitted a total of 10 models, including eight for Andean uplift-dependency vs. two null models where rates are constant through time. We believe that this set of models is lacking in alternative scenarios, and that Andean uplift-dependency could have been selected simply because null constant rate models are unrealistic. Specifically, the study fails to include time-dependent models (see Morlon et al. 2011; Lewitus and Morlon 2018). Time-dependent models represent scenarios of rates changing in time (not constant), but not associated to a specific environmental variable. Thus, here we corrected this analysis in two ways: (i) we separated *Phymaturus* and *Liolaemus* clades for independent analyses, because RPANDA does not consider possible rate shifts along the phylogeny, or potentially different rates across clades; and (ii) we expanded comparisons among a total of 42 models (details in Table S9), including time-dependency, as well as an alternative environmental variable based on global temperature variation during the past 67 mya (from Zachos et al. 2001; 2008). Ambient temperature is an important environmental parameter in reptiles, given that they are ectotherms. These additional models provide a wider range of alternative scenarios beyond simply constant rates, since we incorporated other alternatives to Andean uplift-dependency beyond a simple null constant model. See extended Materials and Methods and Table S9 in Supplementary Material for further details about these analyses.

Given that our empirical results suggest that *Liolaemus* and *Phymaturus* have different diversification drivers (see results), and the fact that RPANDA is not designed to deal with such scenarios, we conducted a simulation study to explore the impact of ignoring such conditions. Here, we wanted to explore the possible outcomes of RPANDA when large trees include clades influenced by different evolutionary drivers. Given that the program is not designed to deal with such scenarios, it is interesting to know whether: (i) the two true models receive the highest AIC weights, (ii) only one true model receives the highest AIC weights, or (iii) other unrelated models are supported by high AIC weights. We simulated a total of 100 replicated trees under a pure birth model for two different clades (C1 and C2), where the C1 speciation rate is exponentially correlated with temperature (lambda = 0.2; alpha = −0.05), and C2 diversification was influenced as follows:

i. C2 with constant speciation rate (lambda = 0.12);
ii. C2 under time-dependent exponentially correlated speciation rate (lambda = 0.2, alpha = −0.025);
iii. C2 under Andean uplift-dependent exponentially correlated speciation rate (lambda = 0.045, alpha = 0.0005); Parameter values used for simulations were selected in order to produce approximately similar tip numbers between C1 and C2 (average ~ 220-250 each). Both C1 and C2 are set to be 40 mya old.

## Results and discussion

See also extended results in Supplementary Materials.

### Disparate patterns of diversification in the lizard family Liolaemidae: further analyses contradict previous study by Esquerré et al. (2019)

#### Heterogeneous diversification within the lizard family Liolaemidae

BAMM estimation of speciation and extinction rates in the Liolaemidae phylogeny (Fig. 1a-b) displays two shifts (PP = 0.4; Table S2), including the origin of the genus *Phymaturus* (red), and the *Liolaemus elongatus* clade (light blue). There are significant differences in speciation and extinction rates between genera (p < 0.001), as clearly shown by the distributions of parameter estimations (Fig. 2). Specifically, the genus *Phymaturus* has the highest speciation rate that is also associated with a high extinction rate. This result is concordant with another study using a different phylogenetic tree (Olave et al. *in press*), and also supported by similar results recovered using ClaDS (Fig. S2). We note that both programs find two shifts to accelerated speciation rates involving clades that were classified entirely as “Andean” by Esquerré et al. (Table 2).

**Figure 1:**
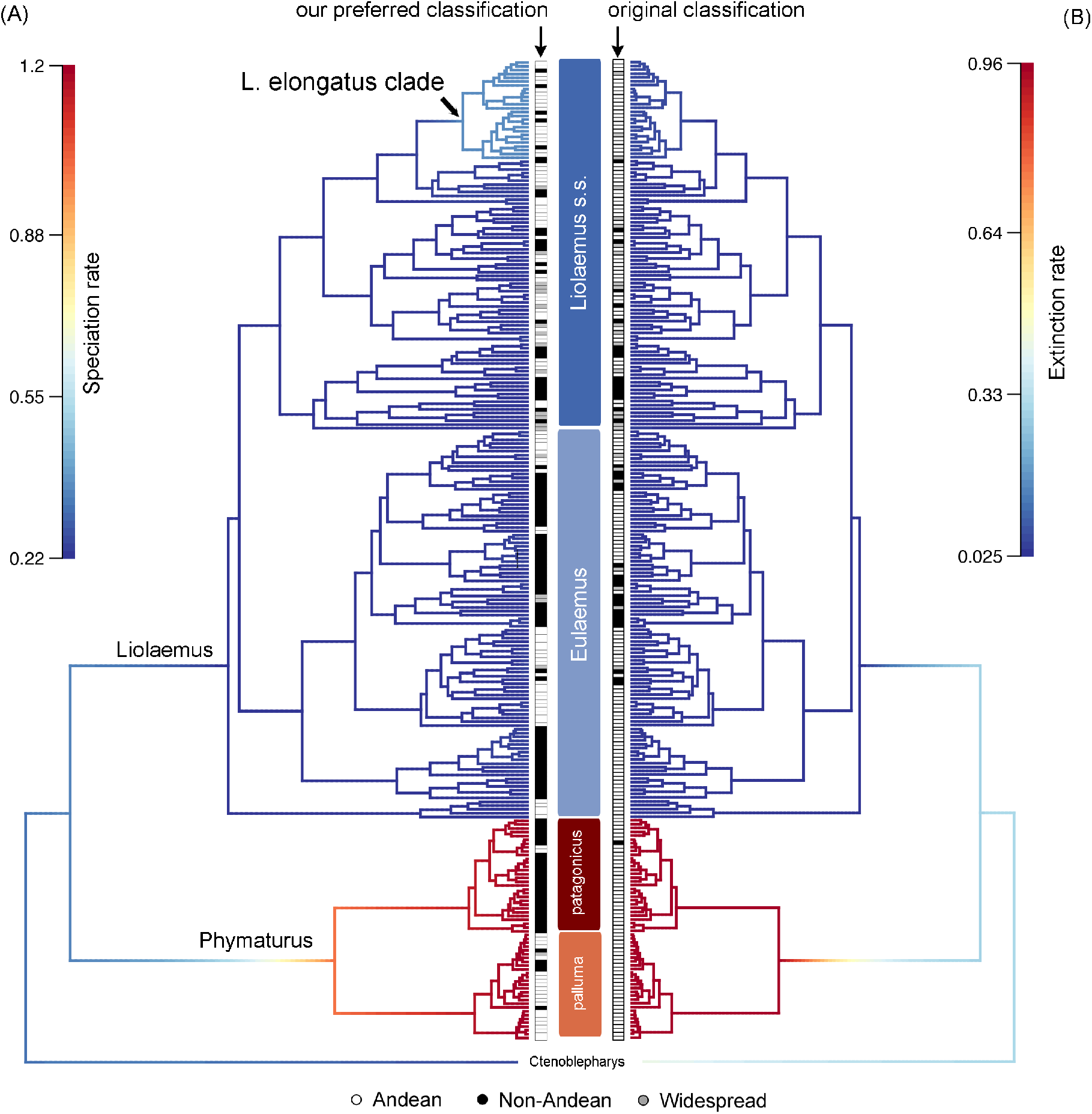
Color-coded phylogenetic trees for the speciation (A) and extinction (B) rates through time for the Liolaemidae inferred by BAMM. Our preferred geographic classification of species is shown on the left tree and the original geographic classification on the right tree; corresponding to Andean, non-Andean, and widespread, shown at each tip in white, black, and gray, respectively.

**Figure 2:**
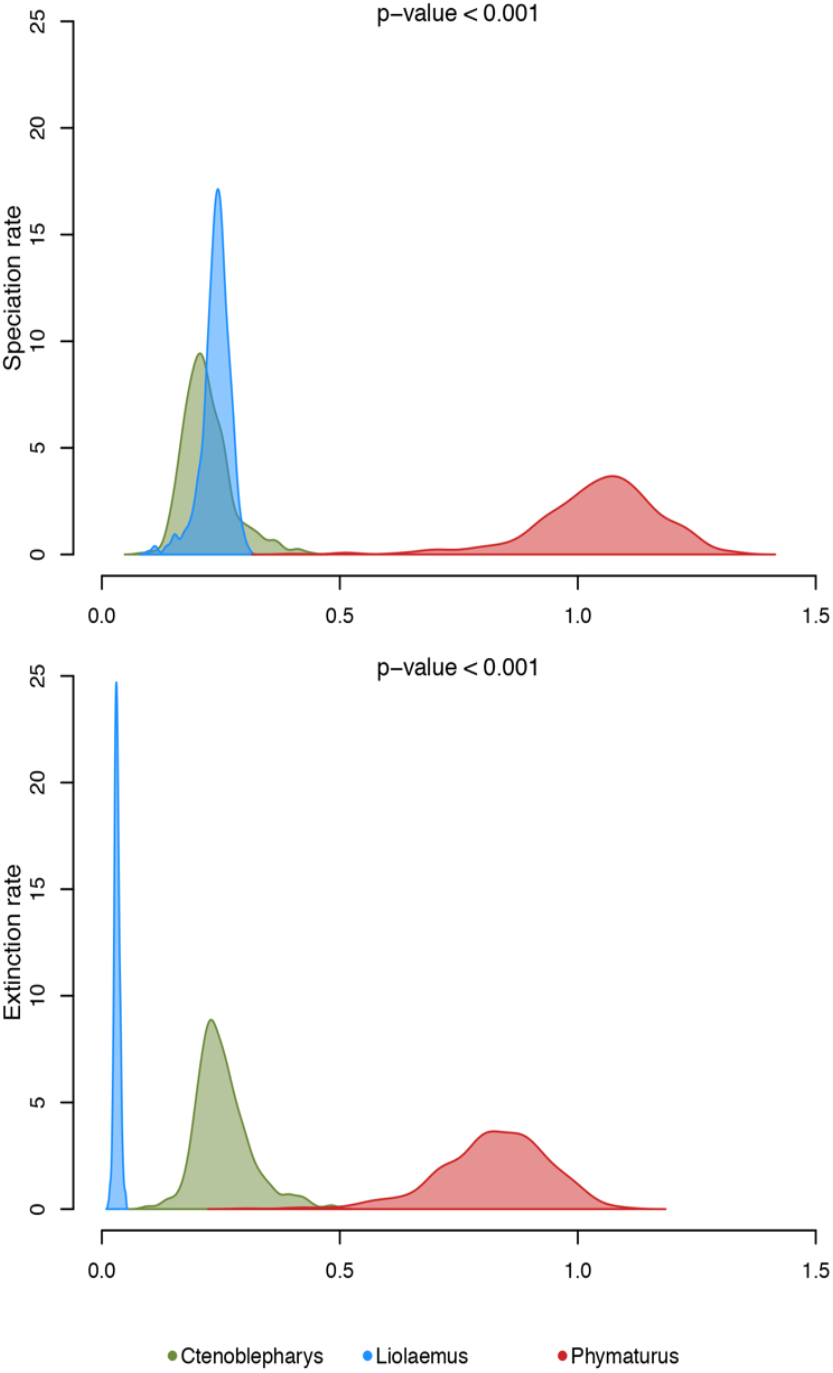
Speciation and extinction rates obtained for *Ctenoblepharys* (green), *Liolaemus* (blue) and *Phymaturus* (red). The density plots are constructed using the mean obtained from each of the last 500 trees of the MCMC runs for phylogenetic estimation.

**Table 2:**
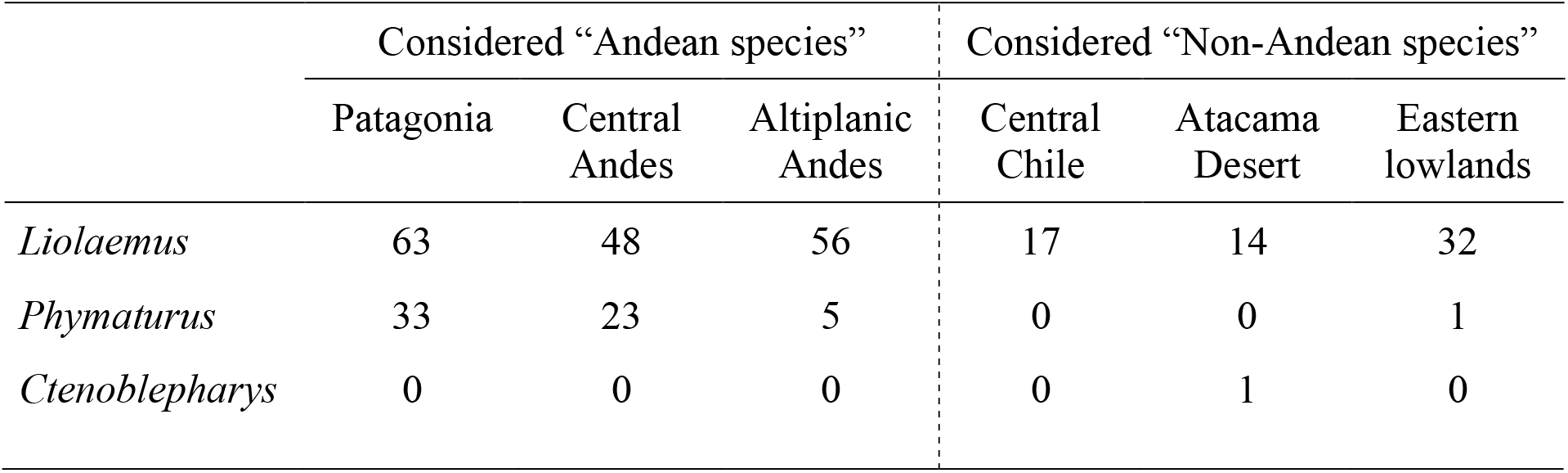
Species counts for the geographic classification from Esquerré et al. (2019); taken from their supplementary material. Note that some species were classified as both Andean and non-Andean as “widespread” (classification details in Table S8).

|  | Considered “Andean species” |  |  | Considered “Non-Andean species” |  |  |
| --- | --- | --- | --- | --- | --- | --- |
|  | Patagonia | Central Andes | Altiplanic Andes | Central Chile | Atacama Desert | Eastern lowlands |
| *Liolaemus* | 63 | 48 | 56 | 17 | 14 | 32 |
| *Phymaturus* | 33 | 23 | 5 | 0 | 0 | 1 |
| *Ctenoblepharys* | 0 | 0 | 0 | 0 | 1 | 0 |

#### Exploration of the relationship between speciation and extinction rates vs. maximum altitude

We constructed linear models between the maximum altitude (MA) of species occurrence records, and the species-specific speciation and extinction rates. Linear models reveal nonsignificant correlations of the speciation/extinction rates with the MA for all target clades (Table S5). Scatter plots show no relationship between rates and MA (Fig. 3; Figs. S4-S10).

**Figure 3:**
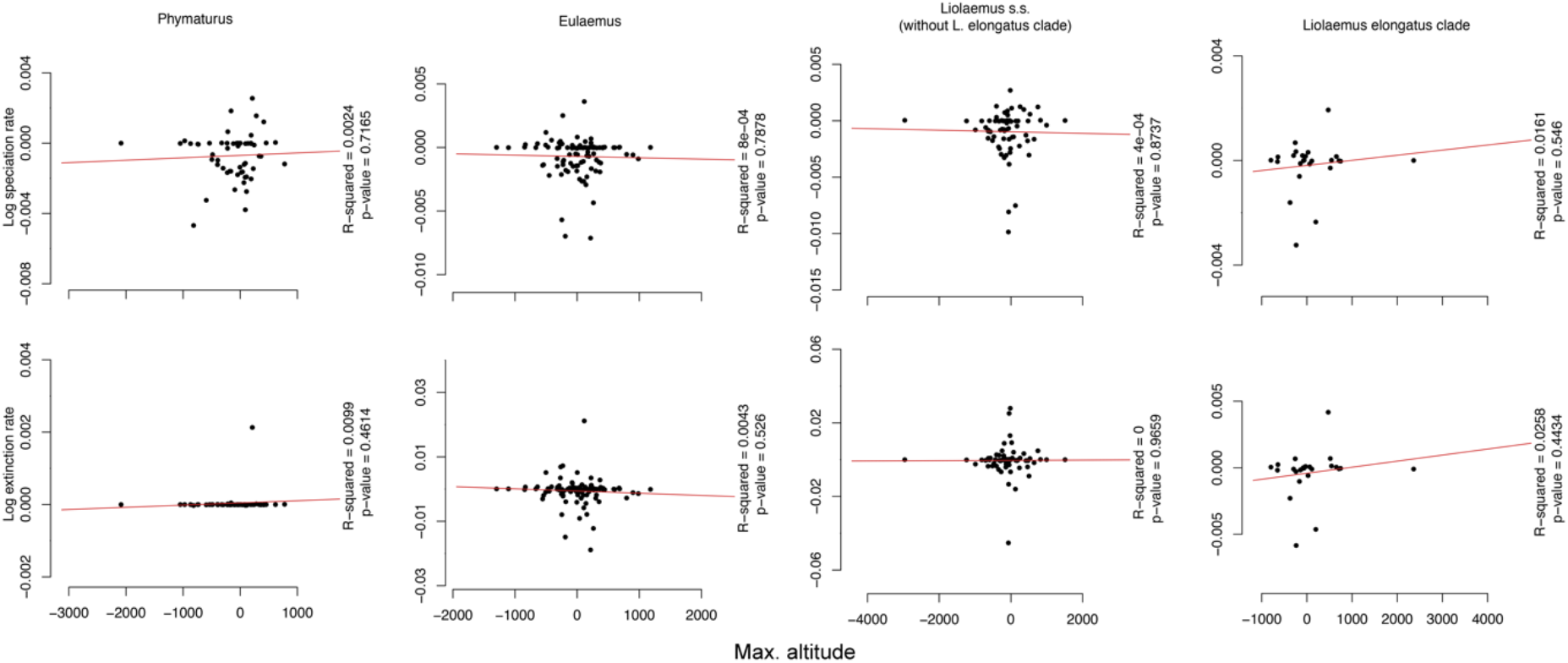
Phylogenetically controlled linear regressions of the log transformed speciation/extinction rates as a function of the maximum altitude of the species occurrence, for different target clades: *Phymaturus* genus, *Eulaemus* subgenus, *Liolaemus* sensu stricto (s.s.) subgenus when excluding the *L. elongatus* clade, and with the *L. elongatus* clade. See also the Figures S3-S9 for more regressions, and Table S5 for full results of the linear models.

#### Incorporating heterogeneous diversification: GeoHiSSE results

Speciation and extinction rate estimates generated by GeoHiSSE are highly congruent with BAMM and ClaDS estimates, with higher rates in *Phymaturus* than *Liolaemus* (Fig. 1, 4 and S2). Note that GeoHiSSE model selection was different when considering the two classifications (Table S7). Following our preferred classification (Fig. 4), the M6 (CID—GeoHiSSE, two hidden rate classes) was associated to the greatest AIC weight (=0.5), followed by M12 (CID—GeoHiSSE extirpation, two hidden rate classes; AIC weight = 0.29). Other CID models showed lower support (M3, M19, M9, M20, M5 and M11; Fig. 4; Table S7). On the other hand, results based on the original geographic classification by Esquerré et al. (Table S7), the M3 (CID—GeoHiSSE, three hidden rate classes) are associated to the highest weight (= 0.47), followed by M18 (CID—anagenetic GeoHiSSE, two hidden rate classes; weight = 0.22) and M19 (CID - GeoHiSSE, four rate classes; weight = 0.17). Other models received little support (weight < 0.05; M4, M5, M9 and M20).

**Figure 4:**
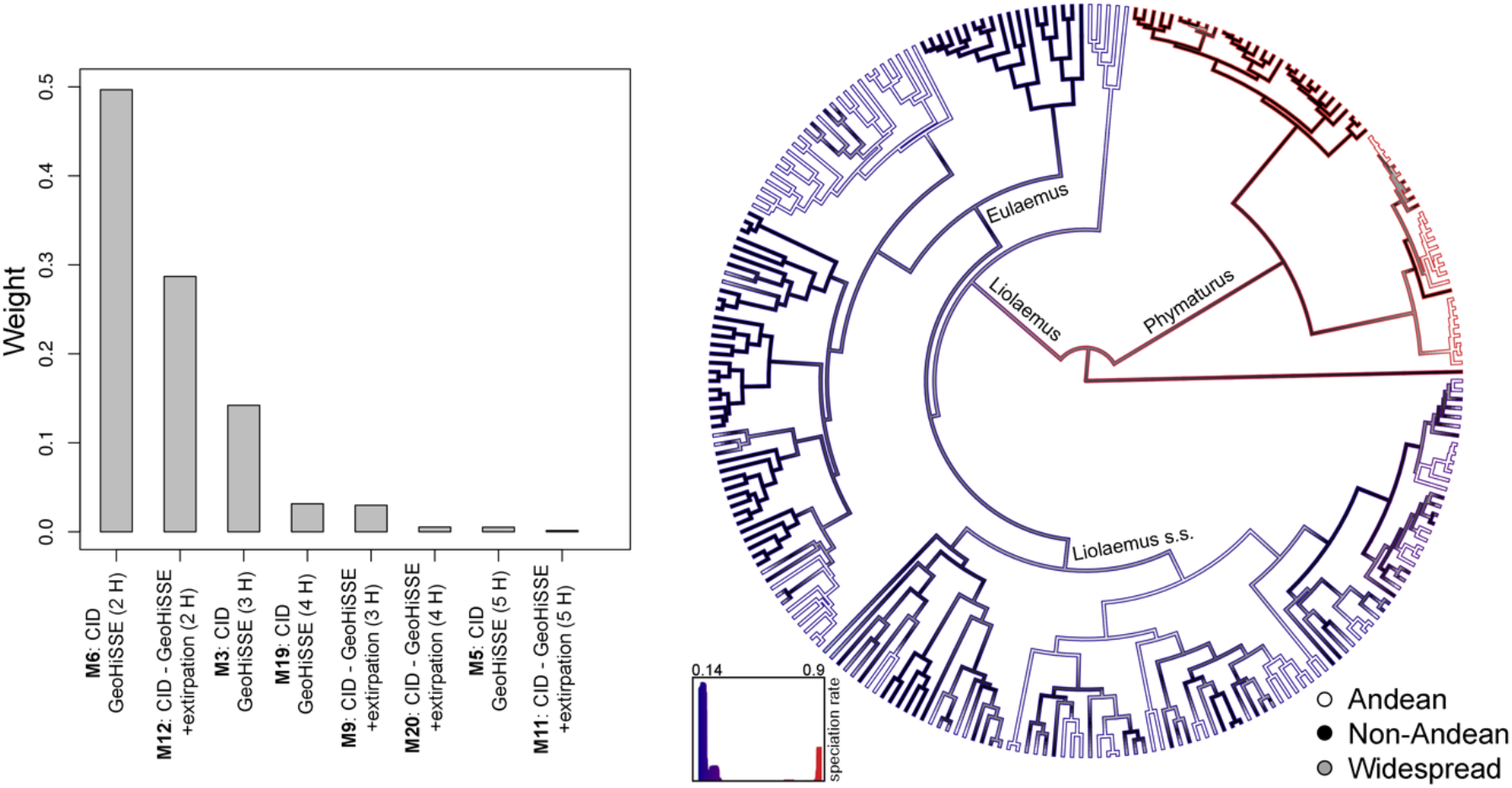
Left, AIC weights of models fitted to the empirical dataset. Right, color-coded phylogenetic tree reflecting the speciation rates (blue-red scale) and geographic-states (whiteblack scale) obtained after averaging parameters of the GeoHiSSE analyses. Results here correspond to those obtained based on our preferred classification. See Table S7 for further result details.

We have demonstrated that the genera *Phymaturus* and *Liolaemus* display clear disparate patterns of diversification with two clear shifts along the phylogeny (Fig. 1, 4 and S2) and that, when incorporating the HiSSE into the models, results show no apparent signal of Andean uplift increasing speciation rates in the Liolaemidae (Fig. 4; Table S7). This is true in both cases, when considering either our preferred geographic classification or the original classification (Table S7). These results show that Esquerré et al. have confounded clade-specific rate accelerations in their distribution-dependent diversification results.

In addition, the GeoHiSSE analyses also return estimates of ancestral distributions (white-black color gradient in phylogenetic tree in Fig. 4). Given the fact that we have changed the geographic classification, these results show that our preferred classification contradicts previous findings by Esquerré et al., using BioGeoBEARS (Matzke 2016) for ancestral reconstructions. This program returned results showing that ancestral lineages in main clades of Liolaemidae were predominantly Andean, leading Esquerré et al. to conclude that the Andes have acted as a “species pump”, and reinforced the idea that Andean uplift has been a key driver in Liolaemidae diversification. In comparison, our preferred classification now shows that the ancestral *Liolaemus* was most likely “widespread” (i.e., both Andean and non-Andean), and that *Phymaturus* is inferred to have a non-Andean ancestor (Fig. 4). The ancestors of the other subgenera are also inferred to be non-Andean or widespread (*Eulaemus:* widespread, *Liolaemus s.s.:* widespread, *P. palluma:* widespread; and *P. patagonicus*: non-Andean; Fig. 4). These results also argue against the Andes acting as “species pump”.

#### Correlative environment-dependent models in RPANDA applied to study the lizard family Liolaemidae

We fitted 42 models to each genus within Liolaemidae using RPANDA. This algorithm does not incorporate “hidden states”, so the only proper way to approach such heterogeneous scenarios is by running separate analyses for each genus (but see extended results in Supplementary Materials and Table S9 for analyses of Liolaemidae as a whole). The best model to describe evolutionary rates in *Liolaemus* is a birth-death model in which speciation and extinction rates are linearly correlated with time (i.e. our model 18; Table S9; Fig. 5A). This model received an AIC weight average of 0.26 ±0.17 (lambda = 0.098, alpha = 0.14; Table S9), followed by a model in which speciation and extinction rates are linearly correlated with global temperature changes (AIC weight mean = 0.08 ± 0.07; model 15; see Table S9 for details of other models that received little support). In contrast, the best model to describe evolutionary rates in *Phymaturus* corresponds to speciation rates exponentially correlated with the interaction between time and global temperature changes (model 19; AIC weight mean = 0.11 ± 0.03; lambda = 1.03, alpha = −0.6; Fig. 5B), followed by another model including an exponential dependency of the interaction between time and global temperature changes (AIC weight mean = 0.06 ± 0.04; model 25). In addition, other models received support and a higher dispersion of estimated AIC weights, reflecting increased uncertainty in *Phymaturus* relative to *Liolaemus*.

**Figure 5:**
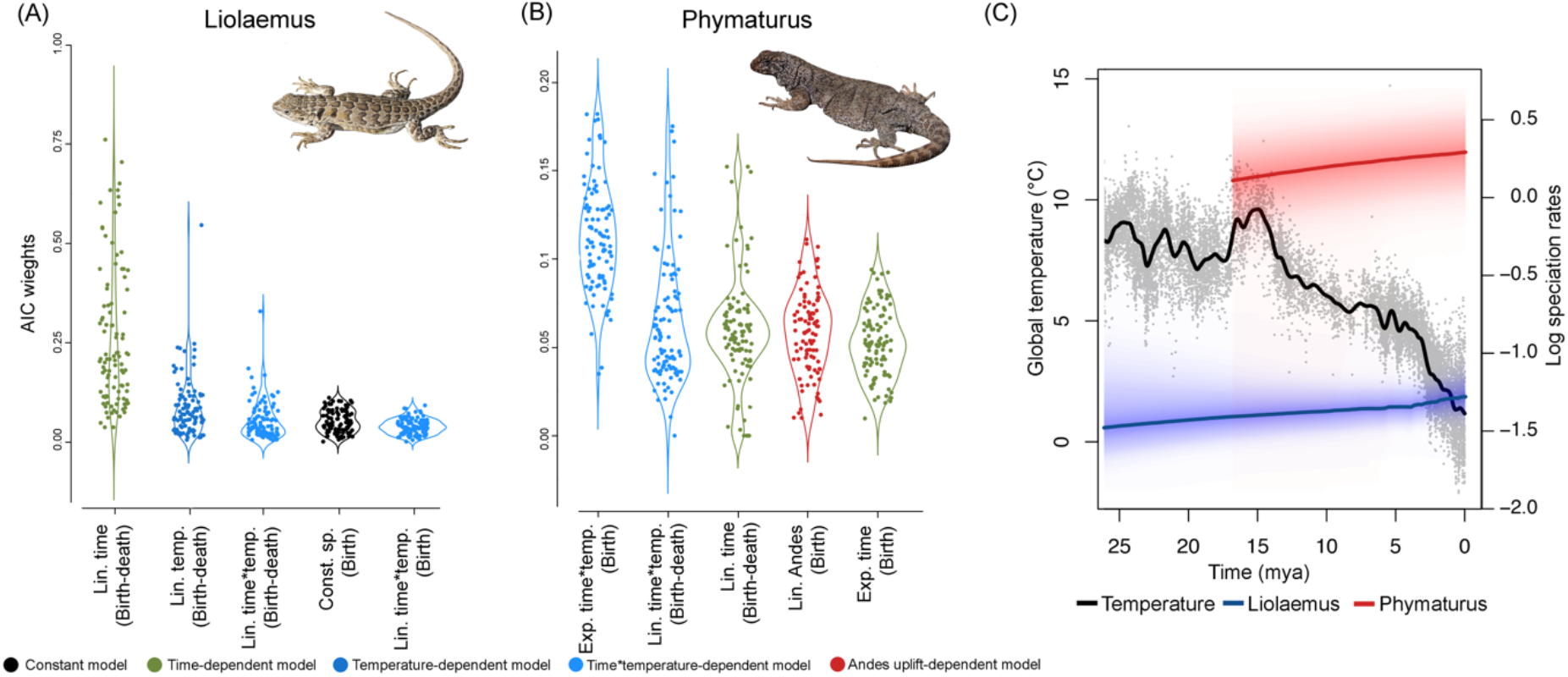
(A-B) Top 5 models (highest AIC weight means) estimated by RPANDA among 42 models for (A) *Liolaemus* genus and (B) *Phymaturus* genus. All detailed results are presented in Table S9. (C) Global temperature changes through time (gray dots) and a smooth spline in black. Speciation rates estimated by BAMM were log transformed for *Liolaemus* (blue) and *Phymaturus* (red).

#### Disparate patterns of diversification in the lizard family Liolaemidae

Interestingly, even though *Liolaemus* has extraordinary species richness (262 described species), our analyses show that its sister group *Phymaturus* evolved under higher speciation rates (44 described species + 23 candidate species included here). Increased speciation rates in *Phymaturus* are best explained by decreasing global temperatures (Fig. 5b). Previous studies have shown that the thermal biology of the cold-adapted *Phymaturus* genus is remarkably similar across species, suggesting that it may be evolutionarily or ecologically constrained (Ibargüengoytía et al. 2008; Cruz et al. 2009). Thus, it is biologically plausible that once the temperature conditions became more favorable for this group, then *Phymaturus’* speciation rates increased (Fig. 5b; c). Different abiotic factors have been suggested to affect South American climates, including Andean uplift as one of the possible promotors (e.g. Gregory-Wodzicki 2000; Blisniuk et al. 2005). Is it possible that the Andes uplift has indirectly driven the increase in speciation rates in *Phymaturus* by promoting local climate change? There has been considerable debate over the role of the Andes relative to South American climate change, including whether the Andes promoted climate change, or global climate change promoted the Andean uplift (Lamb and Davis 2003). However, post-Cenozoic temperature decrease was a global phenomenon (Zachos et al. 2011; 2008), and a combination of relevant factors also acted as locally cooling promotors, such as the Humboldt Current generated by the closure of the Central America Seaway (Hartley 2003; Garreaud et al. 2010). We emphasize that multiple factors have likely driven *Phymaturus* evolution, and assuming a temperature-only dependent diversification is unnecessarily reductionist. We also cannot discard other important aspects in the peculiar biology of this genus. For example, all > 40 cold-adapted species only occur in isolated patches of rock outcrops; therefore, migration between populations is likely severely limited relative to *Liolaemus* (Vicenzi et al. 2017). The strong fidelity of *Phymaturus* species to specific microhabitats, “islands” of big boulders with deep crevices in volcanic cliffs, peaks and plateaus (Cei 1986), might have promoted speciation over short periods of time, as is the case in other lizard species in both wild and urban populations (e.g. Templeton et al. 2001; Thompson et al. 2018).

Specialization in *Phymaturus* has been discussed in other studies (e.g. Reaney et al. 2018; Marín-Gonzalez et al. 2018; Olave et al. *in press*); specialization may be advantageous if it results in efficient selection for adaptation to a stable and narrow niche, reducing the cost of trade-offs by abandoning traits needed to utilize a wider range of resources (Futuyma and Moreno 1988). However, these shorter term microevolutionary benefits may come at the cost of longer term macroevolutionary success (e.g. Agnarsson et al. 2006; Anacker et al. 2011; Forister et al. 2012; Armbruster 2014). Thus, it is possible that the same factors promoting speciation in *Phymaturus* might also explain its high extinction rates as a trade-off (Fig. 1 and 2). The contrasting biology between the sister groups *Liolaemus* and *Phymaturus* (Table 1) makes the Liolaemidae an extraordinarily rich clade in which to explore the cost-benefits of lineages becoming generalists vs. specialists, and their roles shaping macroevolutionary dynamics (Olave et al. *in press*).

### The importance of considering lineage diversification heterogeneity in macroevolutionary studies: warnings in implementation of GeoHiSSE and RPANDA programs

#### Assessing the power of GeoHiSSE models: are hidden states improving state-dependent diversification models?

Previously, it has been shown that including hidden states to the model is a plausible solution to account for diversification heterogeneity and improve the accuracy of the original SSE models (Caetano et al. 2018). We assessed the power of GeoHiSSE analyses by simulating random permutations of geographic areas, and show that most of the CID models received the highest weights over distribution-dependent models (~85%). This is undoubtedly an improvement over the high level of false positives reported for the original SSE models (Rabosky and Goldberg 2015; 2017). However, we still found a relatively high rate of distribution-dependent models receiving the highest weights (= 15%) when considering all 35 models and permutations based on the original geographic classifications (Fig. 6a), and 16% with simulations based on our preferred classification (Table S7). Interestingly, all state-dependent models receiving the highest weights are equal or greater than the maximum number of free parameters (= 15 - 21) in the most complex CID model (M11 and M17). Further, most of these models are supported by high AIC weights (Fig. 6b). We show that excluding models with >15 free parameters (i.e. selection among 32 models) reduced these rates to 7% (Fig. 6c) following the original classification, and to a ~10% rate given the permutations on our preferred classification (Table S7; see also extended results in Supplementary Materials for further details).

**Figure 6:**
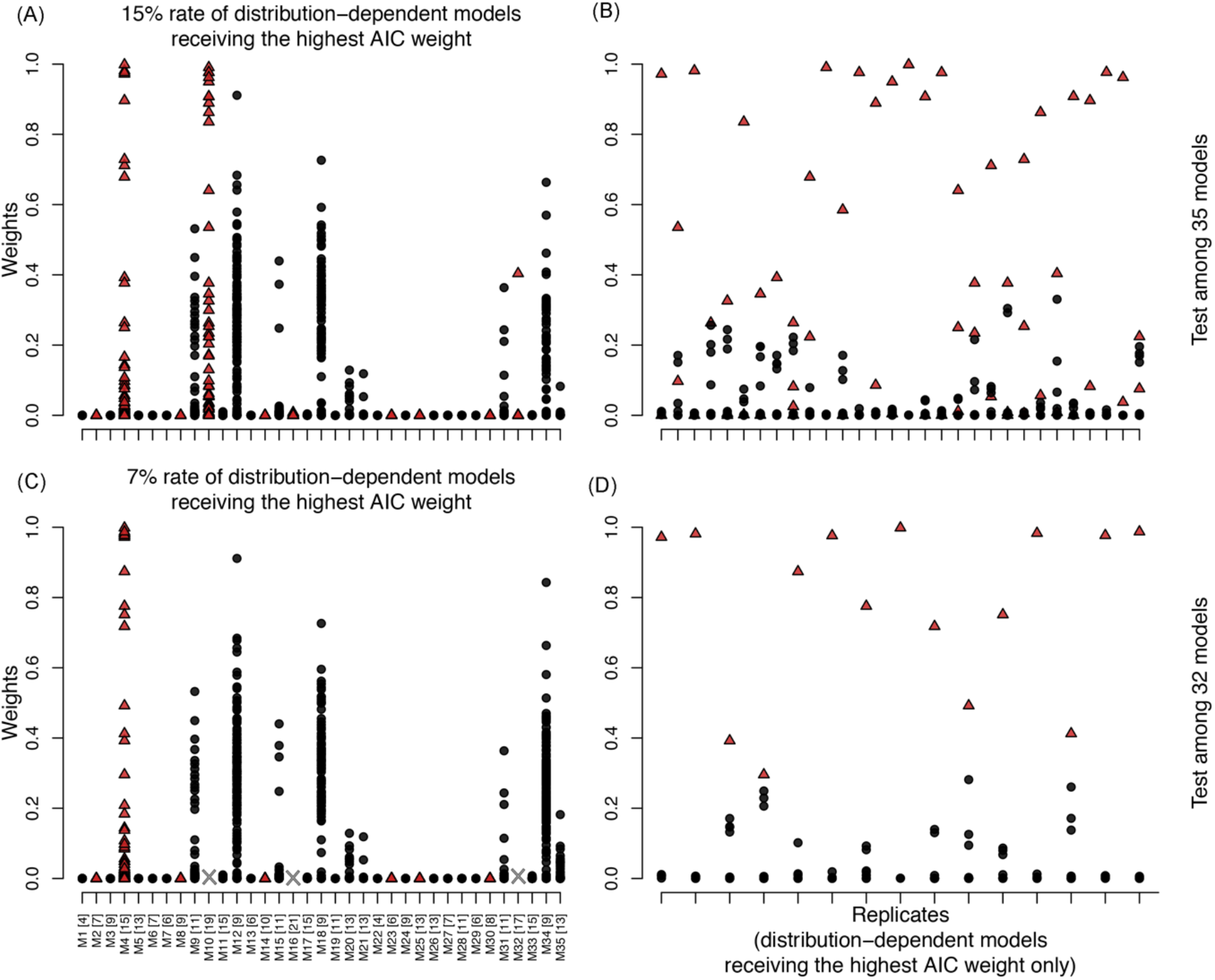
AIC weights among replicated simulations based on the original classification. First row corresponds to analyses performed among 35 models, while the second row corresponds to analyses after removing distribution-dependent models with more than 15 free parameters (32 models total). Left column displays AIC weight per replicate for each model. Right column displays the AIC weight of the different models per each replicate, for which a distribution-dependent model received the highest AIC weights. Red triangles represent geographic-dependent models and black dots geographic independent models (CID). See Table S7 for further result details.

Given the recent documentation that a proper set of state-independent models for HiSSE methods should, necessarily, have the same number of free diversification parameters than the state-dependent models for a “fair” comparison (Caetano et al. 2018), we think that the original set of 35 models should be taken with caution. This issue was paradoxically shown in Caetano et al. (2018) for BiHiSSE implementation, but not corrected among the 35 proposed models for GeoHiSSE in the same paper. Our study does not mean that the same power estimations will necessarily apply to another empirical phylogeny, but we have demonstrated the importance of assessing power before applying HiSSE models, as well as reinforced the notion of potential risks related to the set of models (and number of free parameters) implemented for HiSSE hypothesis tests.

#### Impact of heterogeneous trees in RPANDA: trees including clades following different dependent-drivers affect results

Inspired in our empirical results, we performed a simulation study to address how RPANDA analyses could be affected when two different clades in a phylogenetic tree evolve following different drivers. An extensive simulation study to test the performance of RPANDA was performed by Lewitus and Morlon (2018), including exploration of different proportions of missing tips, accurate model selection, and also rate shifts along the phylogeny. While the authors show that RPANDA seem to be robust when rate shifts are present along the phylogeny under their simulated conditions, here we focused on a different type of heterogeneous scenario.

Our results show that, even though RPANDA can accurately recover the true scenario when there is a single driver for speciation rates in a phylogenetic tree (Fig. S13; Table S10), there is a range of different outcomes when each clade evolves in response to different drivers of diversification, including selecting partially correct models to completely misleading results (Fig. 7). On the one hand, in a scenario where one clade follows a temperature-dependent evolution, while the other clade has constant rates (Fig. 7, left; Table S10), the models with highest AIC weights are two temperature-dependent models (model 4 and 2; model 2 is the true model), followed by a constant rate model (model 1; also a true model). Here, recovering the true models with high support might represent the best possible outcome. However, our second set of simulations included a temperature-dependent + time-dependent trees, and the best-supported models are two unrelated Andean uplift-dependent scenarios (model 2b and 4b; Fig. 7, middle; Table S10). Further, our 3^rd^ set of simulations included trees evolving following two different environmental variables (temperature + Andean uplift; Fig. 7, right; Table S10), and also returns the highest support for an unrelated model (time-dependent; model 3), followed by a temperature-dependent model (model 4; however not the true model).

**Figure 7:**
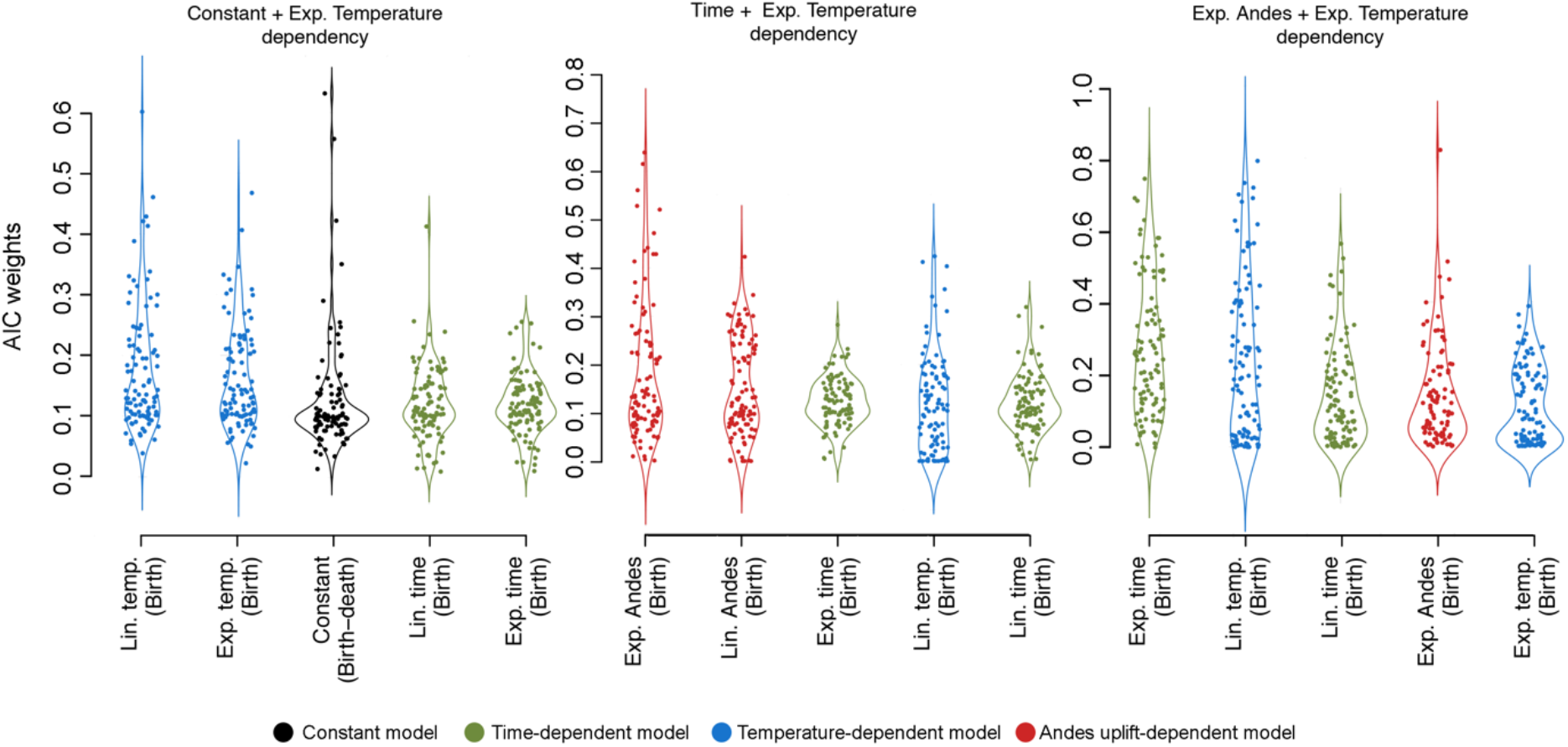
AIC weight obtained for the simulation studies including heterogeneous scenarios analyzed in RPANDA program. Top 5 models (highest AIC weight means) are shown for the different simulated scenarios, including trees comprising one clade (C1) evolving under temperature-dependency plus a second clade (C2) evolving either (i) under constant speciation rates (left), (ii) with speciation rates exponentially correlated with time (middle) and (iii) with speciation rates exponentially correlated with the Andes uplift as a second environmental variable (right). Further results are shown in Table S10.

Our results highlight clear potential risks of ignoring such heterogeneous diversification histories in RPANDA. Currently, the only way to circumvent such an issue is to recognize potential differences *a priori*, and separate clades into different analyses. However, while hidden states are not incorporated in RPANDA, it is clearly difficult in practice to recognize which clades should be separated, since identifying speciation rate shifts might not be sufficient. Specifically, two clades with similar rates might follow different environmental drivers. Thus, recognizing clades with important biological differences is key to assess potential differences *a priori*. As we have shown here, the two contrasting groups of *Liolaemus* and *Phymaturus* not only differ in speciation rates, but also in several aspects of their biology (Table 1). Such biological differences are expected to affect the macroevolutionary patterns (Li et al. 2018). Empirical studies must explore such possibilities when using models that do not allow heterogeneous trees, as is it the case of the current version of RPANDA.

## Conclusions

Incorporating large trees for macroevolutionary studies has the advantage of providing larger datasets, and presumably more power. However, it is important to keep in mind the value of incorporating heterogeneous diversification rates into models for state- and environment-dependent hypothesis testing. We disagree with the main conclusion of Esquerré et al. that the Andes uplift was the main driver of the diversification of these lizards, by showing that: (i) there is no support for increased speciation rate in Andean species (Fig. 4, S7), (ii) a time-dependent model better explains diversification of the genus *Liolaemus* than Andean uplift (Fig. 5; Table S9), and (iii) a model considering the interaction between time and global temperature changes is a better fit for the genus *Phymaturus* than Andean uplift (Fig. 5; Table S9). As a final remark, our preferred classification of species distributions has now changed inferences in ancestral-range reconstructions, and suggests that ancestors of both genera and all four subgenera were non-Andean or widespread (i.e. both Andean and non-Andean; Fig. 4), thus challenging the Andean “species pump” hypothesis. However, we do not attempt to invalidate the whole work presented by Esquerré et al. As mentioned before, they have presented many other relevant findings about the diversification history of Liolaemidae.

Finally, our study calls attention to the importance of considering heterogenous diversification when implementing state- and environment-dependent hypotheses tests. Specifically, we have demonstrated that trees including clades following different dependent-drivers affect RPANDA results (Fig. 7), and the only way to circumvent such problem is to separate clades for independent analyses, given that this method does not account for hidden states. We also provide recommendations for the implementation of GeoHiSSE models to account for fair comparisons among models included in the analysis. We reinforce the previously known case of removing the distribution-dependent over-parameterized models in GeoHiSSE (Fig. 6), to prevent exceeding the number of free parameters in the most complex null model.

We note that, while here we focused on the recently published Liolaemidae phylogeny as a model system, it is very likely that many earlier empirical studies may have limitations similar to those we report here, and these should be re-assessed.

## Supporting information

Supplementary Material: General Description, Extended Materials and Methods, Extended Results, Table S2 and Figures S1-S13

Table S1: Classifications and BAMM estimations

Table S3: Details of ClaDS parameter estimations.

Table S4: Details of results of phylogenetically uncontrolled linear models fitted for different target clades.

Table S5: Details of results of phylogenetically controlled linear models fitted for different target clades.

Table S6: Details of GeoSSE analyses performed for different target clades.

Table S7: Details of GeoHiSSE results for simulations and empirical analyses.

Table S8: The two classifications used for the GeoHiSSE analyses.

Table S9: Details of 42 models and RPANDA results for the empirical phylogeny of Liolaemidae.

Table S10: Details of RPANDA simulation study results.

## Acknowledgments

We thank Richard Ree and two anonymous reviewers for useful comments made on earlier versions of this article. We also thank D. Esquerré for providing clarification on how they performed their analyses, and useful comments made on earlier versions of this manuscript. We thank all members of the Grupo de Herpetología Patagónica (IPEEC-CONICET) for continuing support. Financial support was provided by ANPCYT-FONCYT 1252/2015 (MM), and a postdoctoral fellowship (MO) from the Alexander von Humboldt Foundation at the University of Konstanz, Germany.

## Author contributions

MO designed the study and carried out the analyses. LJA and MM provided recommendations based on the biology of the focal organism. MO, MM and JWS wrote and edited the manuscript. All authors read and approved the final manuscript.

