## Supplementary Material: General Description, Extended Materials and Methods, Extended Results, Table S2 and Figures S1-S13 for "How important is it to consider lineage diversification heterogeneity in macroevolutionary studies: lessons from the lizard family Liolaemidae"

1. **Extended materials and methods (this document)**
2. **Extended results (this document)**
3. **Tables (this document + separate files)**

Table S1 (separate file): Table with details about: (i) species classifications into target clades, (ii) maximum altitude data and (iii) summary statistics (mean, s.d., quartiles) for speciation and extinction rates obtained for the species-specific estimations from BAMM (calculated from 500 trees sampled from the posterior distribution of the MCMC run during the phylogenetic analysis).

Table S2 (this document): Posterior probabilities of estimated shifts in BAMM.

Table S3 (separate file): Details of ClaDS parameter estimations.

Table S4 (separate file): Details of results of phylogenetically uncontrolled linear models fitted for different target clades.

Table S5 (separate file): Details of results of phylogenetically controlled linear models fitted for different target clades.

Table S6 (separate file): Details of GeoSSE analyses performed for different target clades.

Table S7 (separate file): Details of GeoHiSSE results for simulations and empirical analyses.

Table S8 (separate file): The two classifications used for the GeoHiSSE analyses.

Table S9 (separate file): Details of 42 models and RPANDA results for the empirical phylogeny of Liolaemidae.

Table S10 (separate file): Details of RPANDA simulation study results.

4. **Figures (this document)**

Figure S1: Color-coded phylogenetic trees for the net diversification rates (speciation – extinction) through time for the Liolaemidae inferred by BAMM program.

Figure S2: Color-coded phylogenetic trees for the speciation (A) and extinction (B) rates through time for the Liolaemidae inferred by ClaDS program.

Figure S3: Mean (dots) and 95% credibility intervals estimated for *Liolaemus* clades.

Figure S4: Linear regressions (phylogenetically uncorrected) of speciation/extinction rates as a function of the maximum altitude of the occurrence of the genus *Phymaturus*, subgenus *Eulaemus*, *Liolaemus* sensu stricto (without *L.* *elongatus* clade), and the *L. elongatus* clade*.*

Figure S5: Linear regressions (phylogenetically uncorrected) of speciation/extinction rates as a function of the maximum altitude of the occurrence of the species of Liolaemidae*.*

Figure S6: Phylogenetically controlled linear regressions of the log speciation/extinction rates as a function of the maximum altitude of the occurrence of the species of Liolaemidae*.*

Figure S7*:* Linear regressions (phylogenetically uncorrected) for speciation/extinction rates as a function of the maximum altitude for the occurrence of species of the *Phymaturus palluma* and *Phymaturus patagonicus* clades.

Figure S8: Phylogenetically controlled linear regressions for log speciation/extinction rates as a function of the maximum altitude for the occurrence of species of the *Phymaturus palluma* and *Phymaturus patagonicus* clades.

Figure S9: Linear regressions (phylogenetically uncorrected) of speciation/extinction rates as a function of the maximum altitude of the occurrence of the species for the genus *Liolaemus*, and the *sensu stricto* subgenus *Liolaemus* (including *L. elongatus* clade species).

Figure S10: Phylogenetically controlled linear regressions of log speciation/extinction rates as a function of the maximum altitude of the occurrence for species of the genus *Liolaemus*, and the *sensu stricto* subgenus *Liolaemus* (including *L. elongatus* clade species).

Figure S11: GeoSSE results for the different target clades: Liolaemidae, *Liolaemus*, *Liolaemus* (excluding *L. elongatus* clade) and *Eulaemus*. Original classifications were used in this analysis.

Figure S12: Color-coded phylogenetic tree reflecting the extinction rates and geographic-state obtained after averaging parameters of the GeoHiSSE analyses.

Figure S13: Top five models based on average AIC weights obtained from RPANDA analyses.

**Extended Materials and Methods**

*Phylogenetic tree*

We used the posterior set of trees from the BEAST analysis of Esquerré et al. (2019) to generate a maximum clade credibility tree using TreeAnnotator 2.4 (Bouckaert et al. 2014). The tree includes the monotypic *Ctenoblepharys*, 188 described + 11 undescribed species of *Liolaemus*, and 35 described + 23 undescribed species of *Phymaturus* (73% species coverage of all recognized Liolaemidae). The tree was inferred using the third codon position of three mitochondrial loci (cytb, ND2 and ND4) and six nuclear loci (B1D, EXPH5, KIF24, MXRA5, PRLR and PNN). Loci were concatenated and analyzed with BEAST v2.4.7 (Bouckaert et al. 2014). The single known Liolaemidae fossil was used for calibrations (Albino 2008).

*Identifying possible diversification shifts along the tree and exploration of the relationship between speciation rates vs. altitude*

As in similar models, BAMM assumes the given topology is the true phylogenetic tree, so to account for the topological uncertainty and following the manual’s recommendations, we ran all analyses using 500 trees sampled from the posterior distribution of the BEAST analysis of Esquerré et al. (2019). We informed the proportion of missing taxa using *globalSamplingFraction* = 0.73, thus the program accounts for the missing tips (i.e. 73% coverage). Priors were generated using *setBAMMpriors* in BAMMtools (Rabosky et al. 2014), and we used all 500 obtained means for target groups (genera, subgenera, clades and tips) to construct the final distributions used for all downstream comparisons. All BAMM analyses were run for 5 x 10^6^ generations, sampling every 1,000 generations, with 25% burnin, and assessed convergence using effectiveSize function (all parameters >200). We constructed parameter distributions per genus that captured topological uncertainty. We calculated summary statistics using R (mean, standard deviation and quartiles), and compared statistical differences among specific target clades with ANOVA tests using the R function aov().

We extracted species-specific speciation and extinction rates for different target clades, including family, genus (*Phymaturus* and *Liolaemus*), subgenus (*Eulaemus* and *Liolaemus* sensu stricto), clades within *Phymaturus* (*P. palluma* and *P. patagonicus*) and several smaller clades within *Liolaemus* (Table S1). The analyses were performed considering both controlled and uncontrolled phylogenetic relationships. Phylogenetic independent contrasts were performed using the pic() function of the R package ape (Paradis et al. 2018) and the gls() function of the R package nlme (Pinheiro et al. 2012). We also calculated linear models using the R, with the formula: rate ~ “target clade” * “maximum altitude”. Recognizing criticisms of the BAMM program (Moore et al. 2016; Meyer and Wiens 2018; Meyer et al. 2018; however see also: Rabosky et al. 2017; Rabosky 2018a; 2018b), we also incorporated a second program for comparision with BAMM estimations. Specifically, we estimated speciation and extinction rates using the recently introduced ClaDS algorithm (Maliet et al. 2019). This method allows detection of many small shifts along the phylogeny, while BAMM tends to recover larger shifts. We used the R function fit_ClaDS0() of RPANDA package (Morlon et al. 2016) to infer branch-specific speciation rates.

*Geographic State Speciation and Extinction (GeoSSE).*

*Geographic State Speciation and Extinction (GeoSSE).* We implemented the GeoSSE (Geographic State Speciation and Extinction) models (Goldberg et al. 2011) to test the hypothesis that higher speciation rates are associated with the Andean species. Using GeoSSE and the original geographic classification in Esquerré et al. (2019), we ran all analyses for the Liolaemidae as a single clade, and then also for different nested clades. We used ML to estimate the parameters as a starting point for an MCMC chain of 30,000 generations with a 20% burnin. Convergence was assessed by observing likelihood stabilization through iterations. All analyses were performed in the R package *diversitree* (FitzJohn et al. 2012).

*Geographic Hidden-State Speciation and Extinction (GeoHiSSE).*

We evaluated the set of 35 different models proposed by Caetano et al. (2018). We included simulations based on random permutation of geographic areas to assess power of GeoHiSSE program (Caetano et al. 2018) given the empirical phylogeny of Liolaemidae. Random permutations of geographic distributions were performed using the R function sample() without replacement, following Rabosky and Goldberg (2015). We fitted the models to the real data and selected models based on its weight, after exclusion of models with unrealistically high biological parameter estimations. Specifically, there were cases where the parameter estimations reached the maximum bounds (=100; similar issues reported by Miller et al. 2018). In all cases, we used the R function evaluate.models() to parallelize the GeoHiSSE() function of the *hisse* package (Beaulieu and O’Meara 2016). Finally, we averaged the parameter estimation using the MarginReconGeoSSE() function. Best models were selected based on their AIC weights.

*Correlative environmental-dependent models in RPANDA*

*Empirical data.* We expanded the set of models used in Esquerré et al. (10 models) to a total of 42 models (details in Table S9). Note that our 42 models include the same 10 models used in Esquerré et al. (2019), however we also included time-dependent models and interactions between environmental variables and time. We used the same historical Andean elevation data compiled by Lagomarsino et al. (2016). We implemented the function fit_env (Morlon et al. 2016) with the fraction of extant species = 0.73 to a total of 100 trees, randomly sampled from the posterior distributions of the phylogenies inferred by the BEAST program. Details of formulas are provided in Table S9. We computed the AIC weights using the function akaike.weights of the qpcR package (Ritz and Spiess 2008).

*Simulation studies*. We simulated a total of 100 replicated trees under a pure birth model for two different clades (C1 and C2). We implemented the tess.sim.age function (Höhna 2013) for the time-dependent and constant rate simulations, and sim_env_bd function (Morlon et al. 2011; Condamine et al. 2013) for environmental-dependent simulations. We limited simulations to a minimum of 50 tips per clade, and then fitted a total of eight models for each set of simulations to each clade separately. Specifically, we fitted the four true models for individual clade simulations (mentioned above), in addition to four alternative models including linear responses to time and environmental variables, with a null birth-death model (constant): (i) a pure-birth model where speciation is linearly correlated with time; (ii) a pure-birth model where speciation is linearly correlated with global temperature changes; (iii) a pure-birth model where speciation is linearly correlated with Andean altitudinal variation; and (iv) a birth-death model where speciation and extinction are constant. In order to test whether it is possible to recover the true model when analyzing each clade separately, we fitted models independently to each clade. Then, both clades were combined into a single tree using the R function bind.tree() of the ape package (Paradis et al. 2018), setting root to 50 mya (i.e. ~10 my for split between C1 and C2). Finally, we analyzed the large tree combining two clades (C1 + C2).

**Extended results**

*Phylogenetically uncontrolled linear models*

When considering all species of Liolaemidae (258 tips), the phylogenetically uncorrected data show highly significant differences among genera (p < 2^-16^), and no significant effect of MA (p = 0.808), or their interaction (p = 0.207; Table S4A; Fig. S4). Analyses of *Liolaemus* alone (194 tips) show a highly significant subgenus effect (p = 3.10^-8^), but a non-significant MA effect (p = 0.365), or their interaction (p = 0.57; Table S4E). Equivalent results (i.e. no effect of the MA, but significant clade effect) were found for the subgenus *Liolaemus* sensu stricto (s.s.) when including (97 tips), or excluding the *L. elongatus* clade (71 tips), as well for the *Eulaemus* subgenus (97 tips; see Table S4F-G).

We also found a significant negative linear correlation between MA and speciation (p = 1x10^-4^) and extinction (p-value = 0.0019) rates in the subgenus *Eulaemus*, but a poor fit of the model (R-squared < 0.2; Fig. S3). Analyses of *Phymaturus* alone (58 tips) show a positive linear correlation between speciation rate and MA (Fig. S3), but there is also a clear clustering of the *P. patagonicus* and *P. palluma* clades, both detected by the linear model (clades p < 2^-16^) with a non-significant contribution of the MA (p = 0.158), or their interaction (p = 0.769; Table S4B). Finally, we found a significant correlation between speciation and extinction rates for the *Phymaturus* *palluma* clade alone (28 tips; Fig S7 and Table S4C).

*Implementation of the original GeoSSE model*

We repeated all analyses reported by Esquerré et al. (2019) using GeoSSE, and recovered similar results of increased speciation rates associated for Andean species when considering Liolaemidae as a whole (Fig. S11; Table S6). We first tested the entire clade Liolaemidae, and all analyses reject equal speciation rates in Andean and non-Andean regions (p-value = 0.0001735; Fig. S11; Table S6). Thus, the GeoSSE model returns significantly higher speciation rates in the Andean clade (= 0.27) relative to non-Andean species (= 0.11). This result is consistent with previous findings by Esquerré et al. (2019); i.e., high-elevation Andean environments are significantly associated with high speciation rates in the Liolaemidae. This analysis included all *Phymaturus* species as Andean (Table 2 in the main manuscript), which also displayed a speciation rate three times higher than *Liolaemus* (Fig. 2 in the main manuscript). We re-ran these analyses for the *Liolaemus* species only, which returned only a slightly significant p-value = 0.04114 (Fig. S11) for a higher speciation rate in Andean species (0.1883 vs. 0.1225; Table S6). This signal disappears with the removal of the *L. elongatus* clade (which was classified entirely as Andean; p-value = 0.16863), or when running the test with the *Eulaemus* subgenus alone (p-value = 0.182782; Fig. S11). Thus, when partitioning the analyses, we show that this signal is clearly due to the increased speciation rates in *Phymaturus* and the *L. elongatus* clades, whereas this Andean signal is lost when removing these groups from the analysis (Fig. S11). Thus, the signals of accelerated speciation rates associated with the Andean uplift in the distribution-dependent diversification test using GeoSSE in Esquerré et al. are biased (Fig. S11), due to the two diversification rate shifts along the tree (Fig. 1 of the main article).

*Simulations study to assess power of GeoHiSSE models given the Liolaemidae phylogeny*

The most frequent model (~37%) receiving the highest AIC weights in our simulation study is the M18 (CID—anagenetic GeoHiSSE, 2 hidden rate classes), followed by the M12 (CID—GeoHiSSE + extirpation, 2 hidden rate classes; 30% frequency; see Table S7 for full results). A distribution-dependent model received the highest weights in a total of 15% of the replicates when considering all 35 models and permutations based on the original geographic classifications (Fig. 4a of the main article), and 16% with simulations based on our corrected classification (Table S7). Distribution-dependent models selected correspond to M4 (2 hidden classes; 15 free parameters), M10 (2 hidden classes + extirpation; 19 free parameters), M16 (2 hidden classes [anagenetic]; 21 free parameters), and M32 (2 hidden classes + extirpation [anagenetic]; 17 free parameters). Thus, all selected models have equal or greater number of free parameters than the most complex CID model (M11 and M17). We also computed the rate of parameter estimations reaching the maximum bound (= 100) among models in the simulation study (Table S7), that could indicate cases of unrealistic parameter estimations. In addition, we checked the frequency of these cases affecting the selected models among the false positive cases. We found that only one replicate involved parameter estimations reaching the maximum bounds, specifically our replicate 61 of permutations performed under the original classification. All other replicates selected models with parameter estimation below this bound.

Additionally, although there is clear uncertainty reflected in the weights, most of these models are also associated with high weights (Fig. 4b of the main article). Excluding models with >15 free parameters (i.e. selection among 32 models) reduced these rate to 7% (Fig. 4c of the main article) following the original classification, and to a ~10% rate given the permutations on our corrected classification (Table S7).

Finally, note that for the empirical case of the Liolaemidae phylogeny, nine and five models were removed from the selection due to unrealistic estimates (Table S7; following original classification and our corrected classification, respectively). Specifically, unrealistically high values were inferred. Note that this includes all the models with >15 free parameters, consequently these were removed for weighting and averaging.

*Correlative environmental-dependent models in RPANDA applied to study the Liolaemidae lizard family*

We also ran RPANDA analyses when considering Liolaemidae as a whole (Table S9). This represent a violation to RPANDA, and we included this result only for comparative proposes. We found a time-dependent model better explains the full Liolaemidae tree (model 5; AIC weight mean = 0.15 ± 0.07). The second and third best models correspond to another time-dependent model (model 3; AIC weight mean = 0.11 ± 0.02), and an Andean uplift-dependent model (model 2b; AIC weight mean = 0.11 ± 0.02).

**Supplementary tables**

| Shifts | Posterior probability |
| --- | --- |
| 1 | 0.0940 |
| 2 | 0.4000 |
| 3 | 0.2800 |
| 4 | 0.1400 |
| 5 | 0.0580 |
| 6 | 0.0190 |
| 7 | 0.0059 |
| 8 | 0.0013 |

Table S2: Posterior probabilities of shift estimations by BAMM (analyzed 3751 posterior samples, after 25% burnin).

**Supplementary Figures**

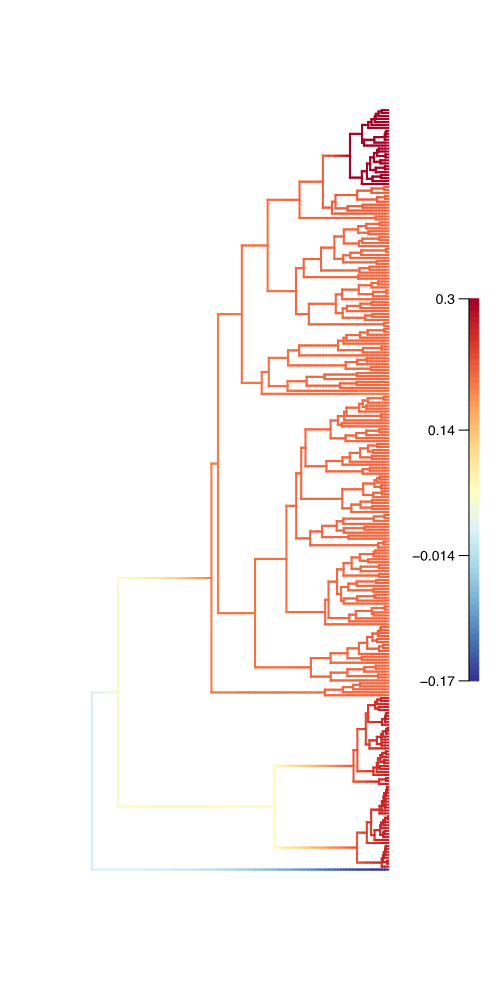

Figure S1: Color-coded phylogenetic trees for the net diversification rates (speciation – extinction) through time for the Liolaemidae inferred by BAMM.

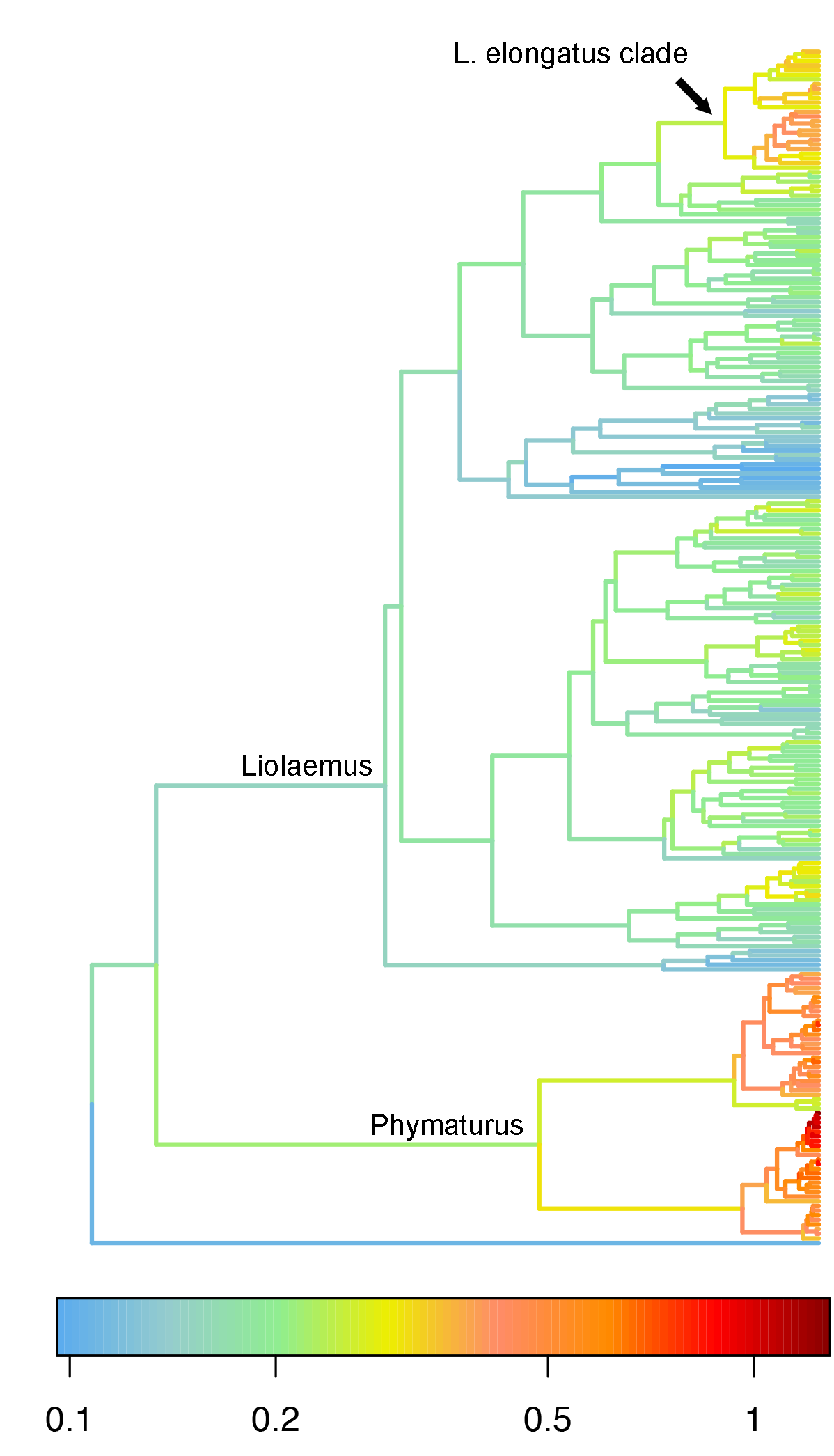

Figure S2: Color-coded phylogenetic trees for the speciation rates through time for the Liolaemidae inferred by ClaDS.

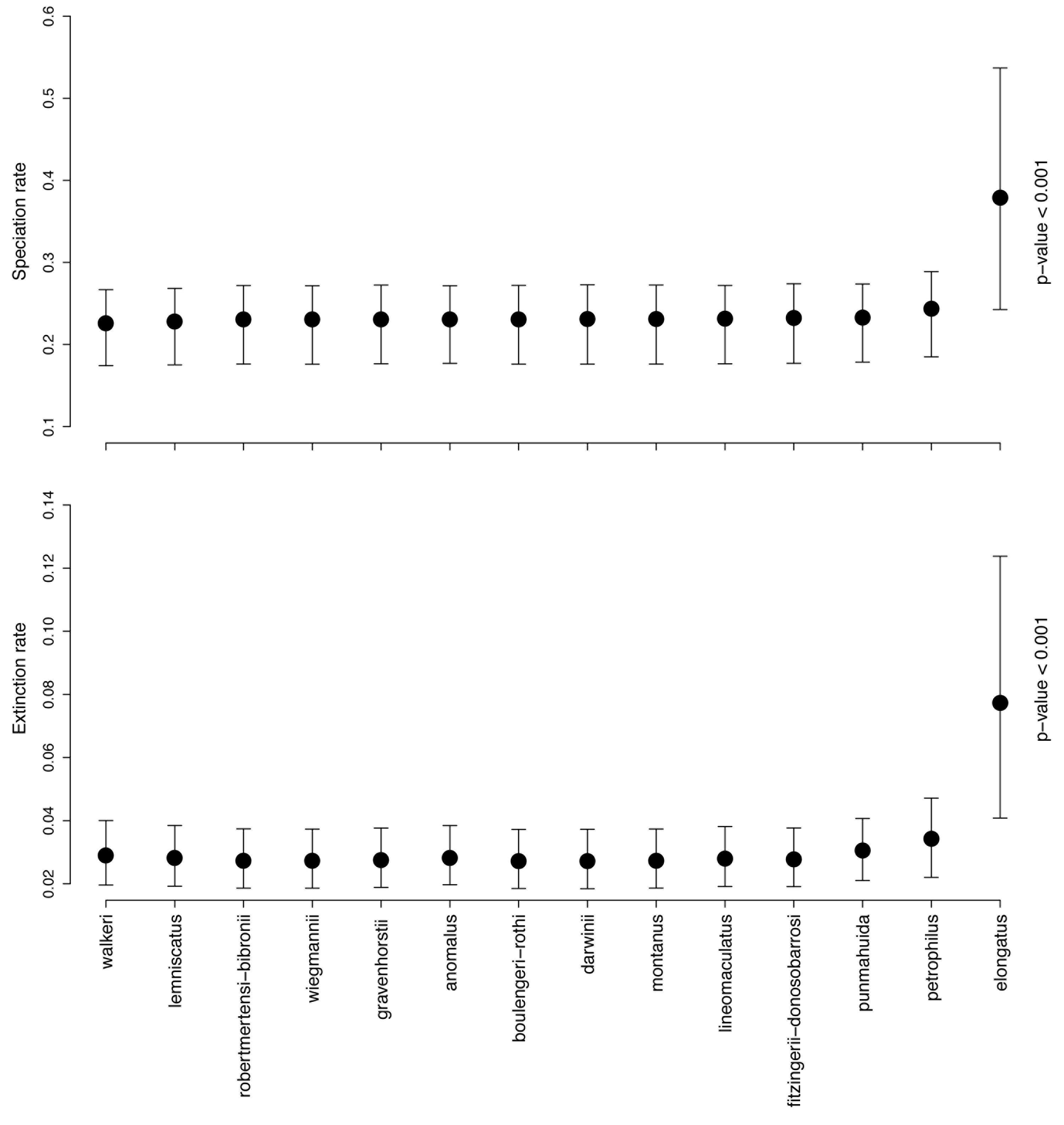

Figure S3: Means (dots) and 95% credibility intervals of speciation and extinction rates estimated for *Liolaemus* clades.

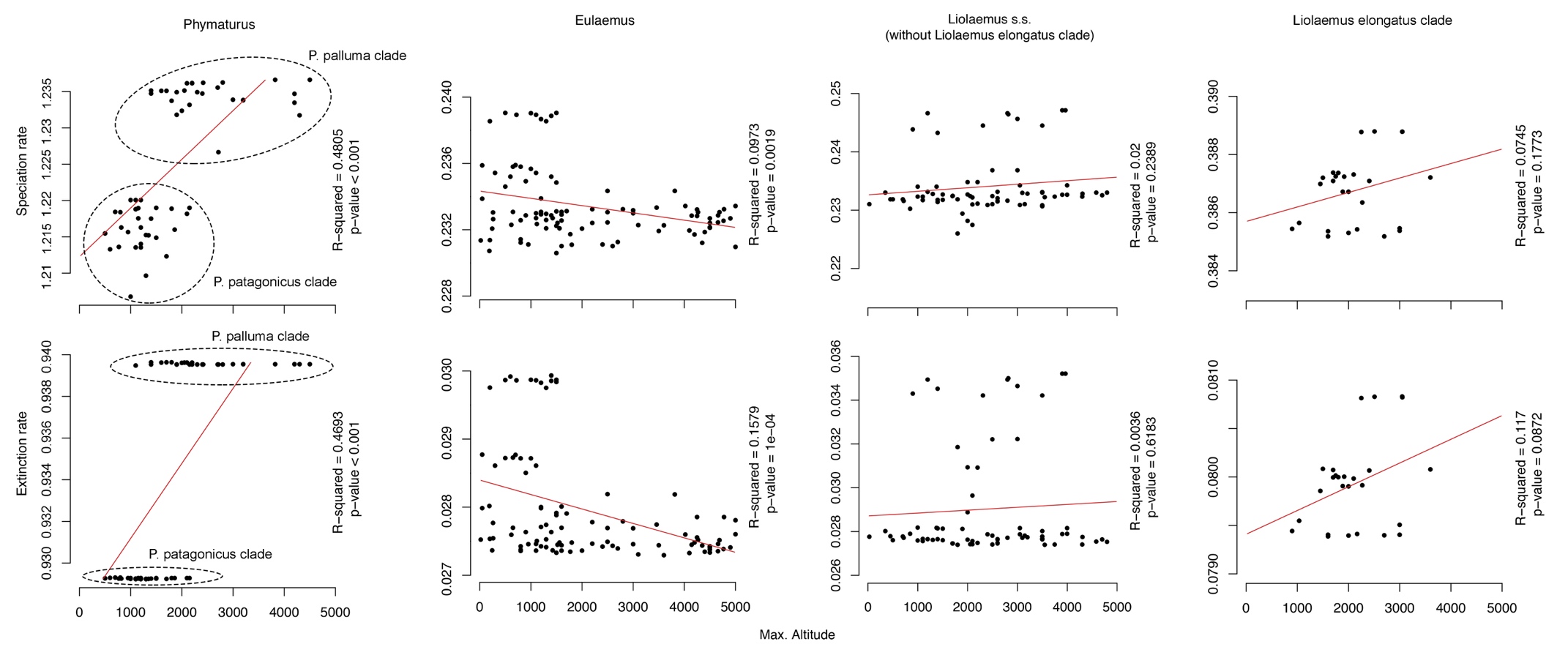

Figure S4: Phylogenetically uncontrolled linear regressions of the speciation/extinction rates as a function of the maximum altitude (meters) of the species occurrences for different target clades: *Phymaturus* genus, *Eulaemus* subgenus, *Liolaemus* sensu strict (s.s.) subgenus when excluding the *L. elongatus* clade, and with the *L. elongatus* clade. See also Table S4 for full results of the phylogenetically uncontrolled linear models.

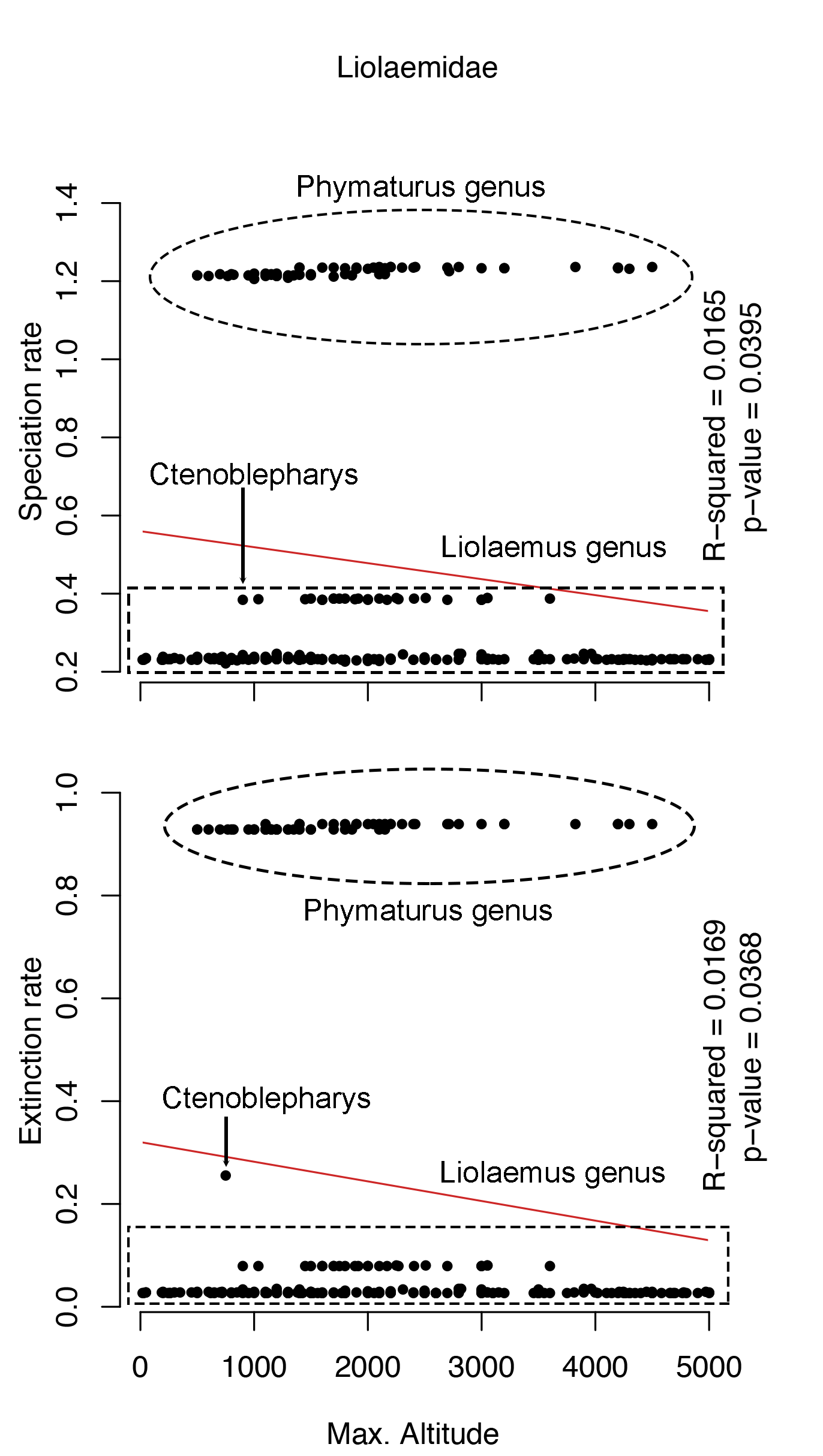

Figure S5: Phylogenetically uncontrolled linear regressions of speciation/extinction rates as a function of the maximum altitude of the occurrence of the species in the family Liolaemidae (see also Table S4A).

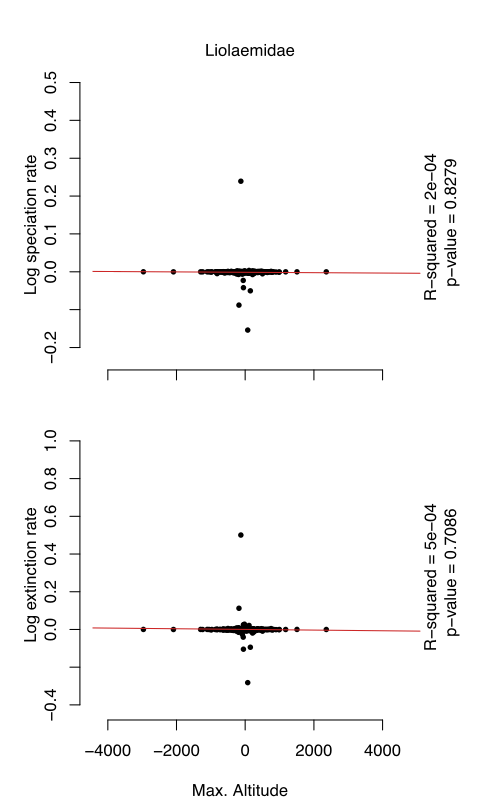

Figure S6: Phylogenetically controlled linear regressions of log speciation/extinction rates as a function of the maximum altitude of the Liolaemidae species*.* When considering all species (258 tips), the regression shows a non-significant correlation between variables when controlling for phylogenetic relationships (both p >0.85; see also Table S5A).

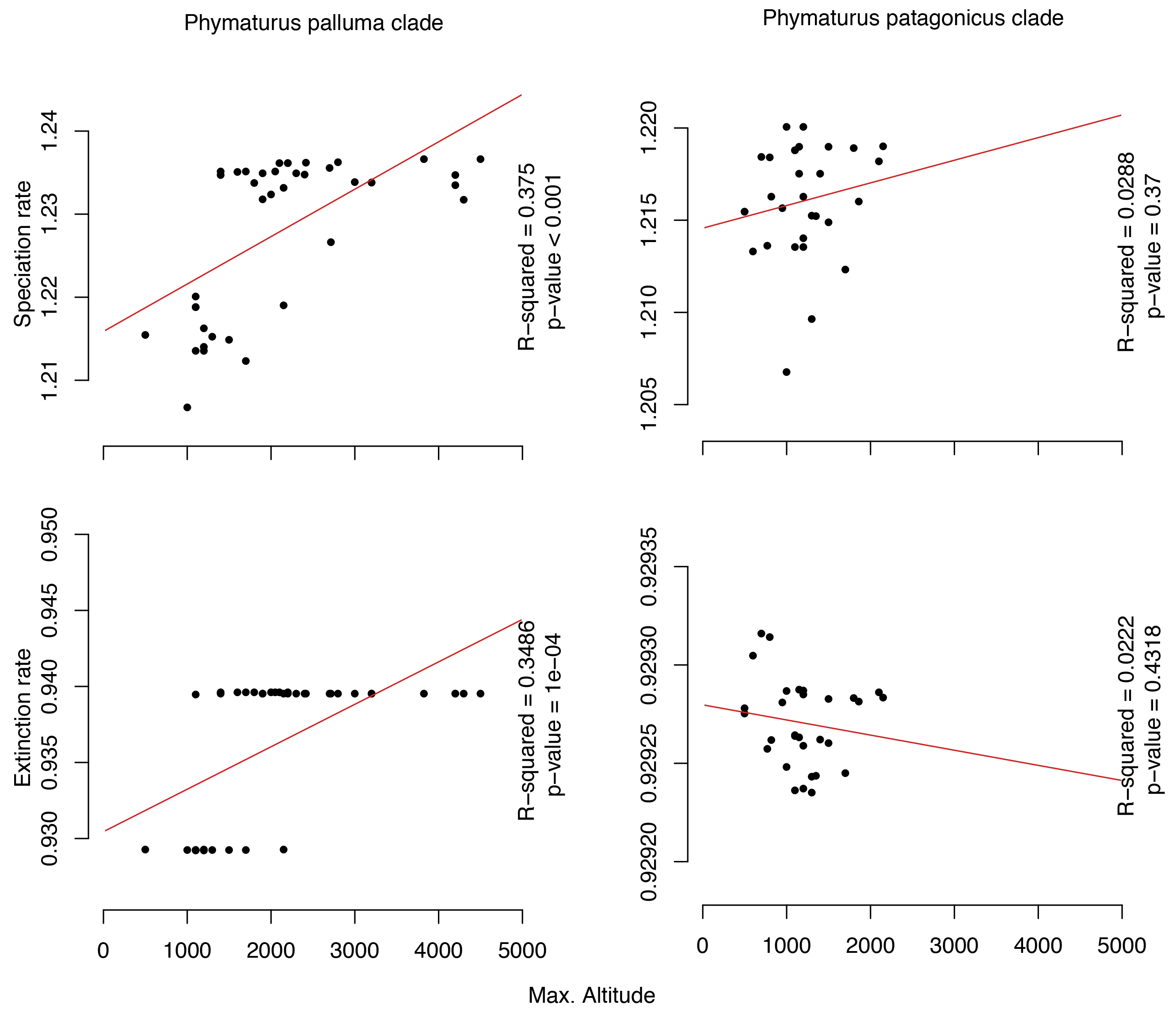

Figure S7: Phylogenetically uncontrolled linear regressions of speciation/extinction rates as a function of the maximum altitude for the species in the *Phymaturus palluma* and *Phymaturus patagonicus* clades (see also Table S4C-D).

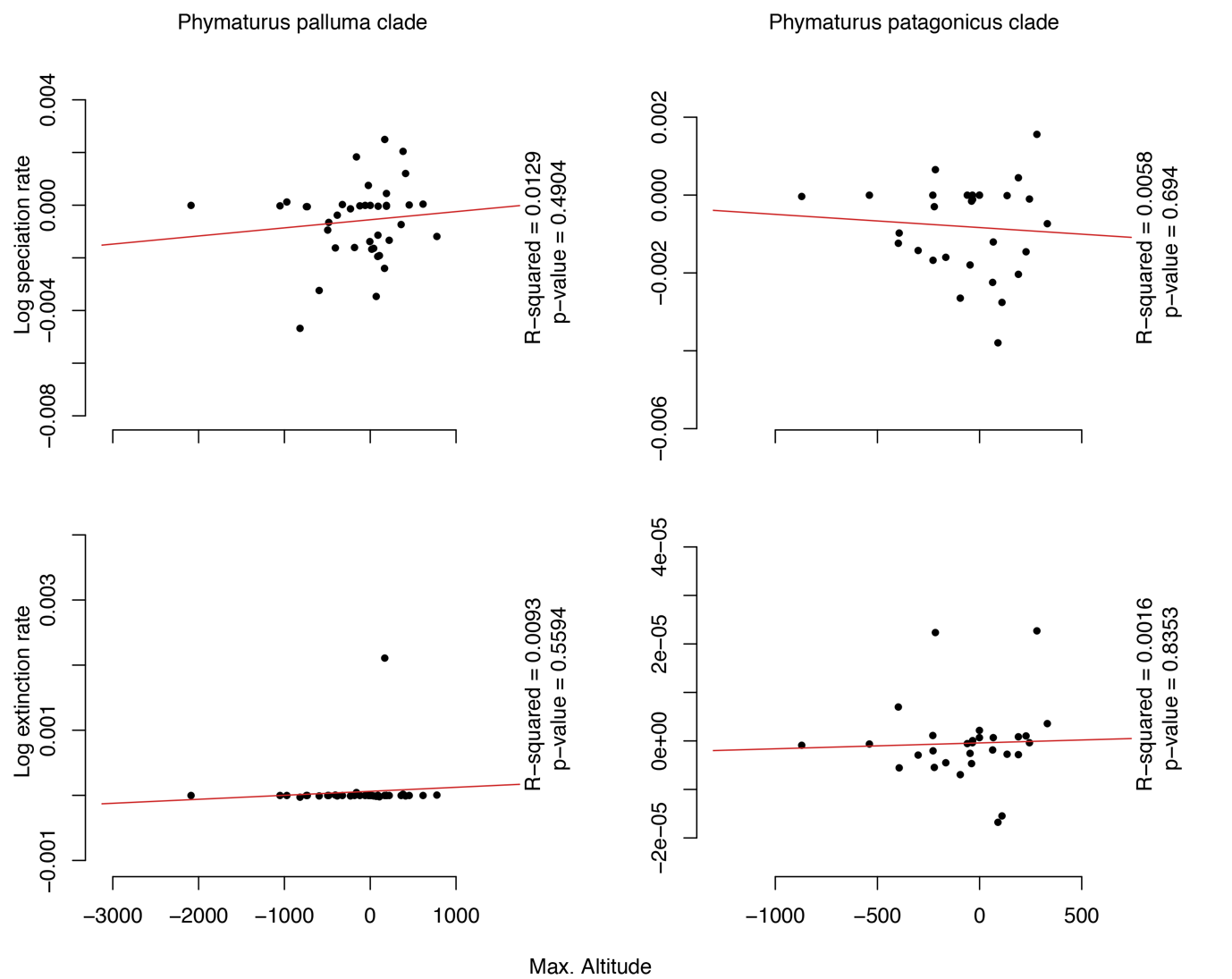

Figure S8: Phylogenetically controlled linear regressions of log speciation/extinction rates as a function of the maximum altitude for species in the *Phymaturus palluma* and *Phymaturus patagonicus* clades (see also Table S5C-D).

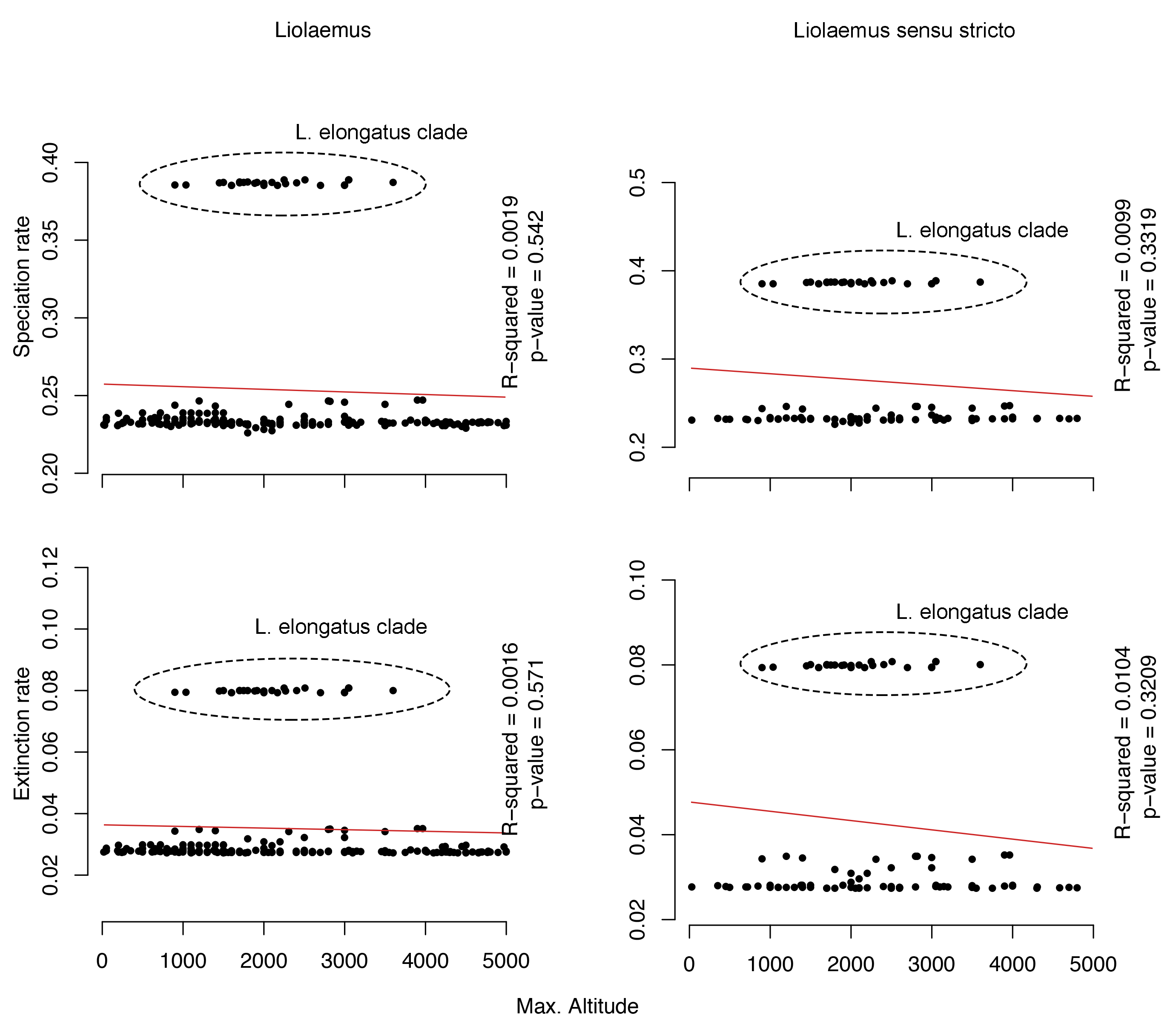

Figure S9: Phylogenetically uncontrolled linear regressions of speciation/extinction rates as a function of the maximum altitude for species of the genus *Liolaemus,* and the subgenus *Liolaemus sensu stricto* (including the *L. elongatus* clade). See also the Table S4E and G.

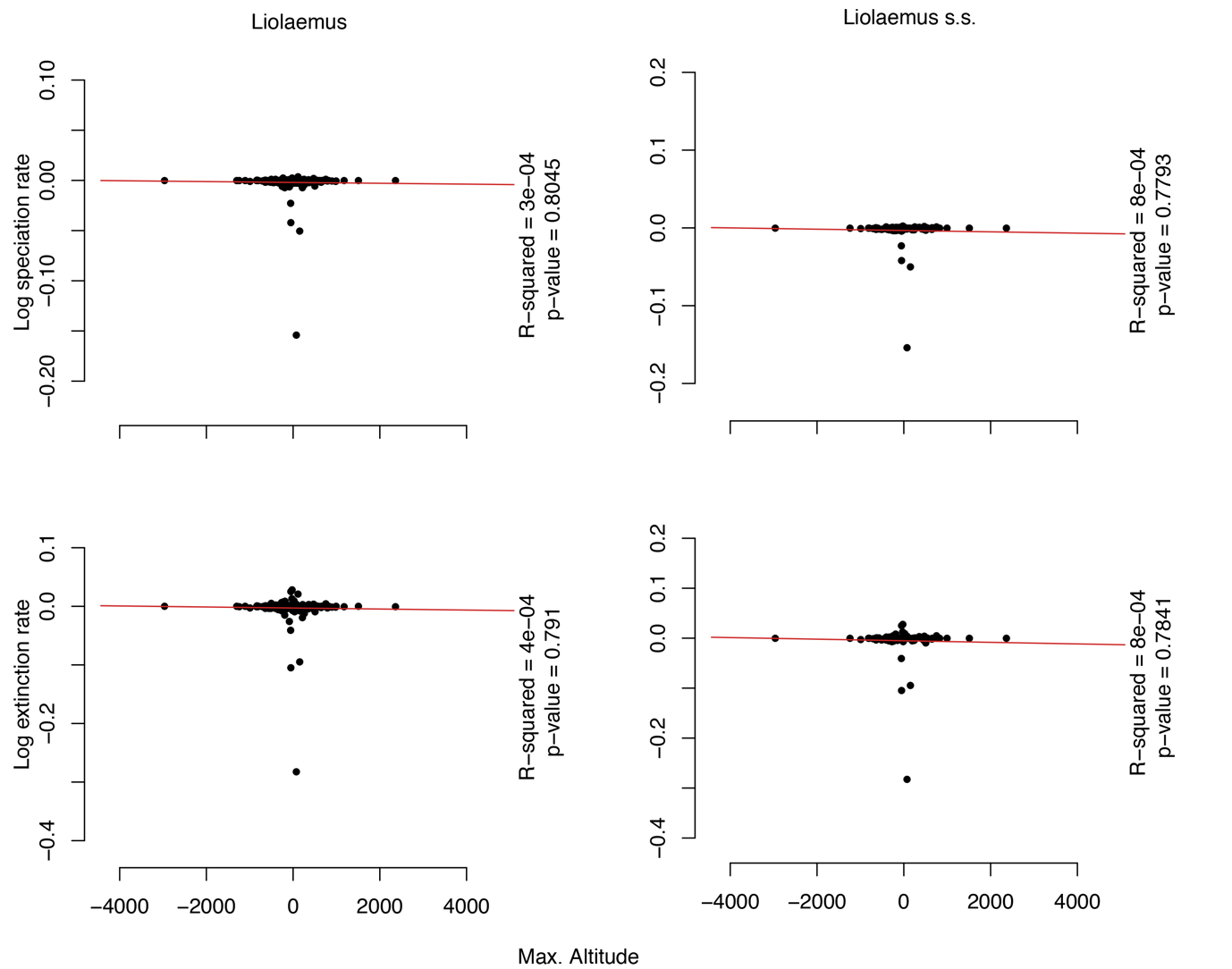

Figure S10: Phylogenetically controlled linear regressions of the log speciation/extinction rates as a function of the maximum altitude for species of the genus *Liolaemus,* and the subgenus *Liolaemus sensu stricto* (including the *L. elongatus* clade species). See also the Table S5E and G.

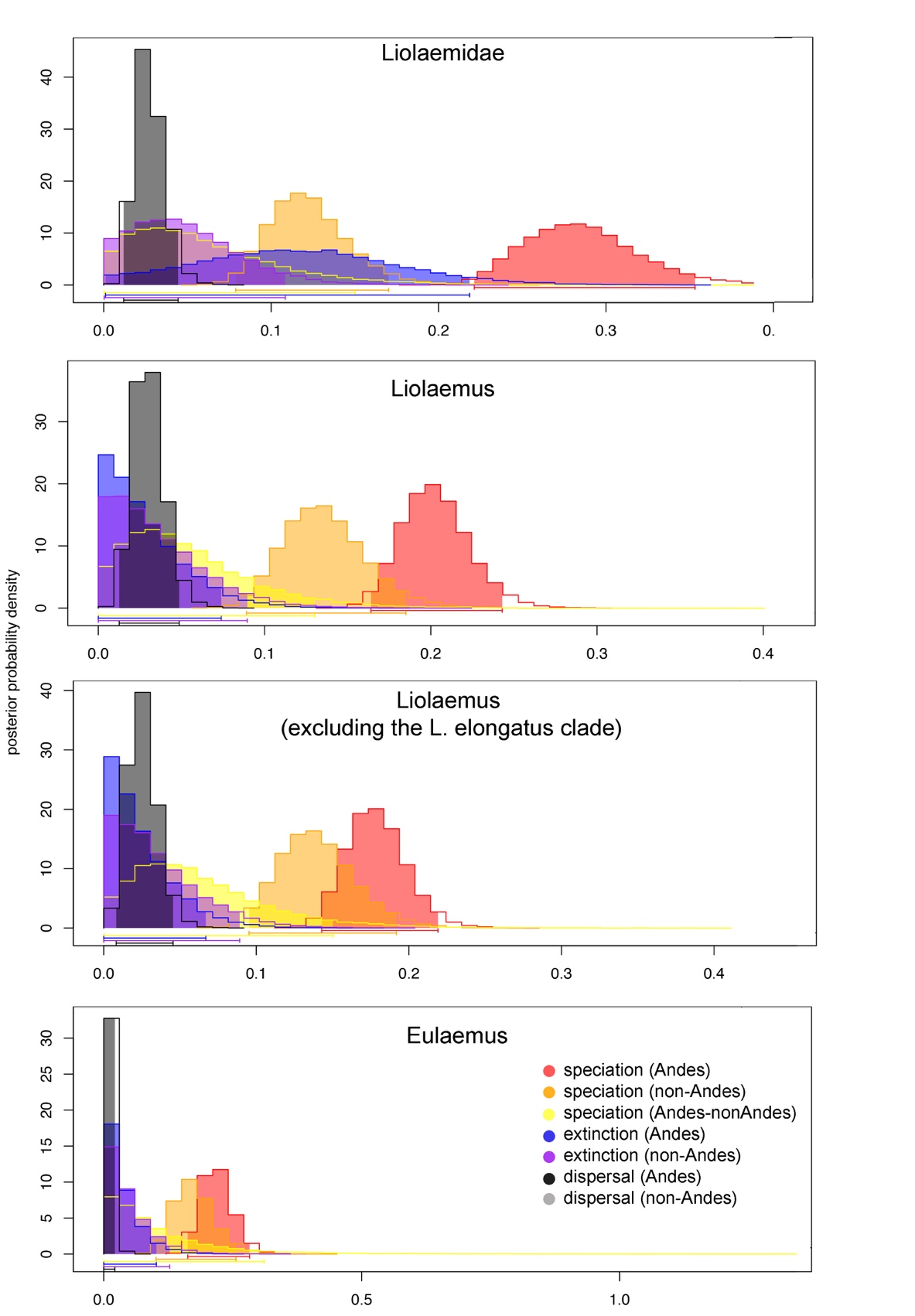

Figure S11: GeoSSE results for all target clades: Liolaemidae, *Liolaemus*, *Liolaemus* (excluding *L. elongatus* clade) and *Eulaemus*. Original classifications were used in this analysis; see also Table S6 for more details.

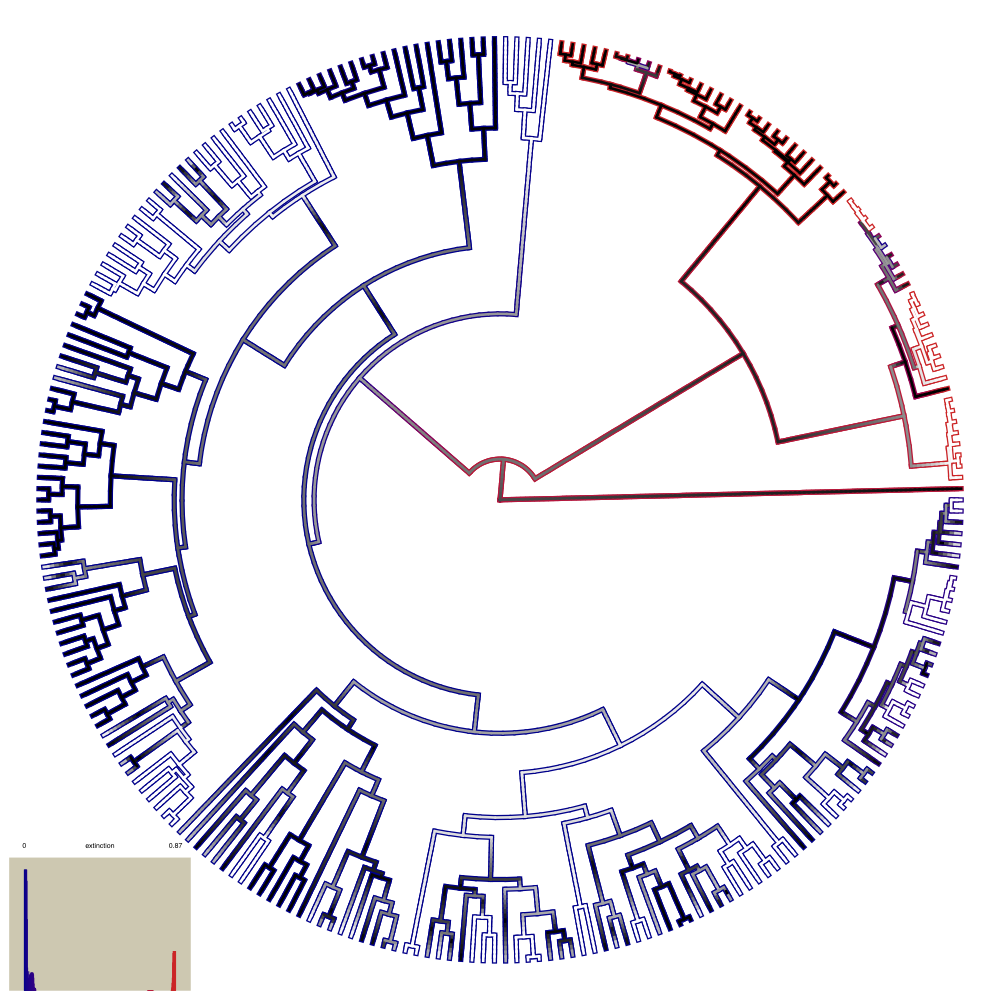

Figure S12: Color-coded phylogenetic tree reflecting the extinction rates (blue-red scale) and geographic-state (based on our corrected classification, Andean: white; non-Andean: black; widespread: gray) obtained after averaging parameters of the GeoHiSSE analyses. See also Table S7.

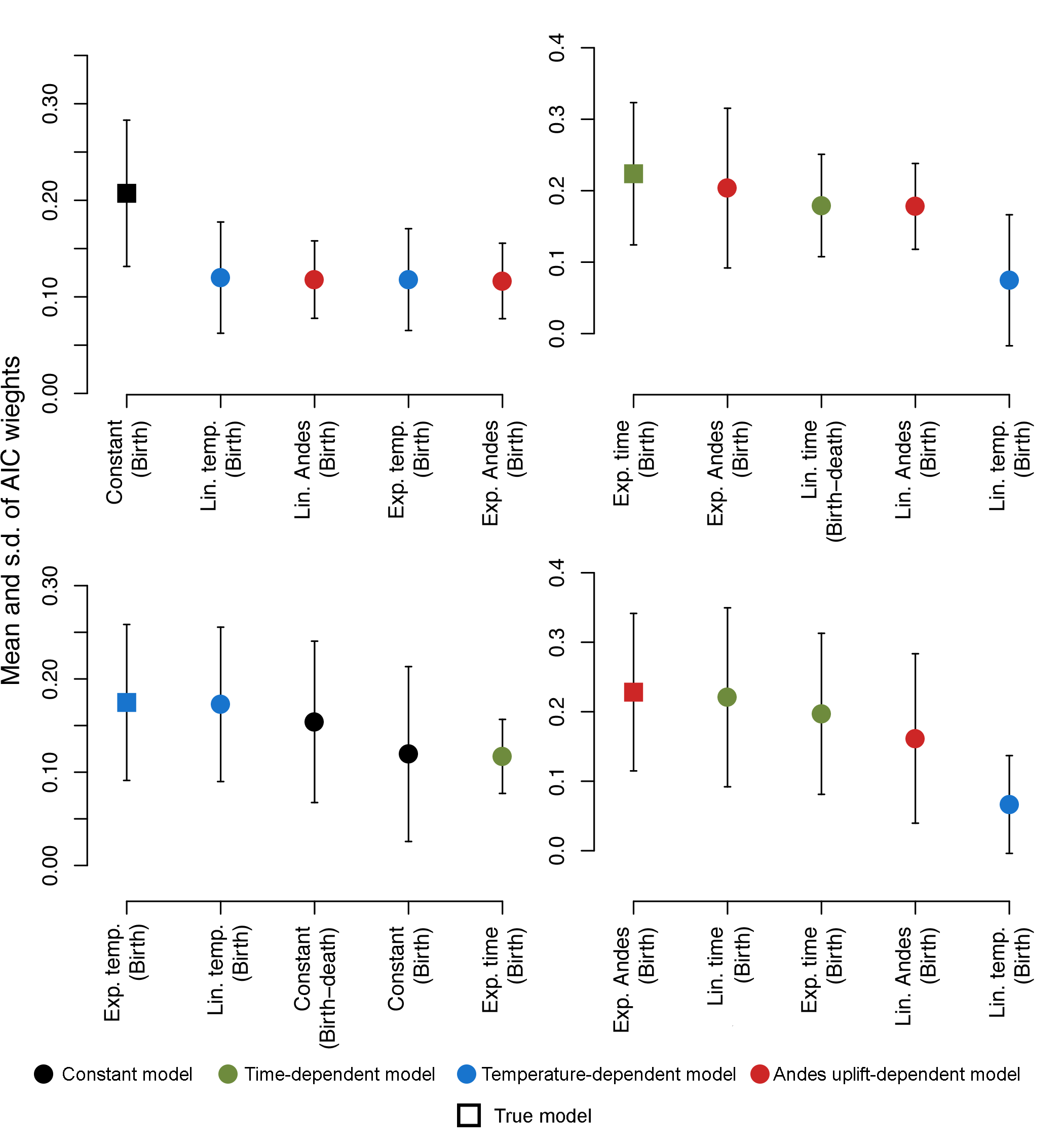

Figure S13: Top five models selected based on average AIC weights obtained from RPANDA analyses based on 100 simulated trees under different models: Pure-birth constant speciation rates through time; Pure-birth model with speciation rates exponentially correlated with time; Pure-birth model with speciation rates exponentially correlated with temperature; Pure-birth model with speciation rates exponentially correlated with Andean uplift. True models used in simulations are represented with a square. See more details in Table S10.
